# Mouse V(D)J Humanization Recapitulates human-like Severe SARS-CoV-2 Immune Imprinting

**DOI:** 10.64898/2025.12.18.694353

**Authors:** Xiao Niu, Fanchong Jian, Yutong Li, Ke Li, Siyu Lei, Weiliang Song, Ruoxi Kong, Xingan Cai, Ran An, Yao Wang, Yifei Huang, Lingling Yu, Wenjing Wang, Haiyan Sun, Yuanling Yu, Jing Wang, Fei Shao, Sai Luo, Yunlong Cao

## Abstract

The mechanisms driving divergent SARS-CoV-2 immune imprinting in populations with distinct SARS-CoV-2 exposure histories remain unclear. However, conventional wild-type mouse models fail to recapitulate severe imprinting, even after potent ancestral-strain mRNA immunization, hindering imprinting-related mechanistic investigation and vaccine evaluation. Here, we surprisingly found that V(D)J-humanized mice could faithfully recapitulate human severe imprinting phenotypes. Comprehensive antibody repertoire analysis and deep mutational scanning-based epitope mapping of 583 monoclonal antibodies from these models revealed that the pre-existing IGHV3-53/66 antibody abundance determine imprinting severity through antibody-mediated masking of Omicron-specific epitopes. Both passive IGHV3-53/66 antibody transfer and IGHV3-53 knock-in were sufficient to induce severe imprinting in wild-type mice. Together, these findings demonstrate that the V(D)J germline repertoire—even a single germline-encoded antibody response—can profoundly shape humoral imprinting severity. Accordingly, we established an IGHV3-53 knock-in mouse model that accurately recapitulates the human antibody landscape, providing a valuable tool for guiding future COVID-19 vaccine updates.

## Introduction

Extensive studies have investigated immune imprinting in SARS-CoV-2 ^1^, yet the reasons why different vaccine platforms produce markedly different imprinting severity have remained unclear. Recipients of mRNA vaccines exhibit pronounced immune imprinting, persistently recalling ancestral Wuhan-Hu-1 (Wuhan) spike-reactive antibodies even upon repeated Omicron exposures, significantly limiting the development of Omicron-specific neutralizing responses ^2–11^. In contrast, individuals only receiving inactivated vaccines and unvaccinated children demonstrate greater adaptability in humoral responses, developing robust Omicron-specific antibodies upon repeated infections or boosters, thus overriding Wuhan immune imprinting ^11–15^.

This distinct imprinting pattern has become increasingly consequential, as it aligns with the emerging global divergence in SARS-CoV-2 epidemiology ^11,16–21^: NB.1.8.1 emerged earlier in 2025 and initially gained a global first-mover advantage, yet XFG subsequently overtook it and became dominant in regions with high mRNA vaccine coverage. By contrast, NB.1.8.1 has remained predominant in countries where inactivated vaccines were widely used, particularly China. A different form of population specificity has also been observed for BA.3.2.2, a highly divergent emerging lineage that is preferentially enriched in weakly imprinted, unvaccinated children.

Understanding the mechanistic basis of this imprinting divergence is critical, not only because it may have directly influenced real-world viral epidemiology—potentially necessitating region-specific vaccine update strategies—but also because these distinct imprinting phenotypes provide a unique opportunity to uncover general principles of humoral immune imprinting applicable to other pathogens, such as influenza.

However, a major obstacle to answering this question is the lack of an animal model that faithfully reproduces severe human-like imprinting while allowing controlled exposure histories and causal perturbation. Conventional wild-type mouse models fail to recapitulate the strong imprinting observed in humans ^2,12,14^, severely hindering mechanistic studies and preclinical evaluation of SARS-CoV-2 vaccines.

In this paper, using V(D)J-humanized mice, we demonstrate faithful recapitulation of strong human-like imprinting and identify pre-existing human IGHV3-53/66-encoded antibodies as the key driver of imprinting severity via epitope masking. Finally, we establish a human IGHV3-53 knock-in mice model that accurately recapitulate human SARS-CoV-2 antibody responses, offering a practical tool for evaluating future vaccine update strategies.

## Results

### V(D)J-humanized mice mimic strong imprinting

Previous population-level studies have shown that SARS-CoV-2 immune imprinting severity is associated with distinct IGHV gene usage patterns ^11,22^. Strongly imprinted individuals, including mRNA-vaccinated individuals, are enriched for IGHV3-53/66 compared with weakly imprinted or unvaccinated populations. Moreover, wild-type mice immunized with ancestral-strain mRNA vaccines fail to recapitulate the strong SARS-CoV-2 immune imprinting observed in the human mRNA-vaccinated cohort ^2^. These observations suggest that the human germline V(D)J repertoire, rather than exposure history alone, may be required for severe SARS-CoV-2 imprinting. Therefore, we employed V(D)J-humanized mice for comparative analysis. These BALB/c-derived mice, commonly used in therapeutic antibody discovery, carry human immunoglobulin V(D)J genes for all heavy chains and kappa light chains (Supplementary Information Figure 4A). Both V(D)J-humanized and wild-type mice were primed with three doses of ancestral mRNA vaccine (encoding the spike protein of SARS-CoV-2 ancestral strain), followed by two booster doses of BA.5 mRNA vaccine, thereby creating a stringent setting to test whether human germline V(D)J genes enable sustained ancestral-strain recall despite Omicron boosting (Figure 1A). Blood, spleens, and lymph nodes were collected one week after the third ancestral dose and each BA.5 dose for serological and flow cytometry analyses.

**Figure 1.**
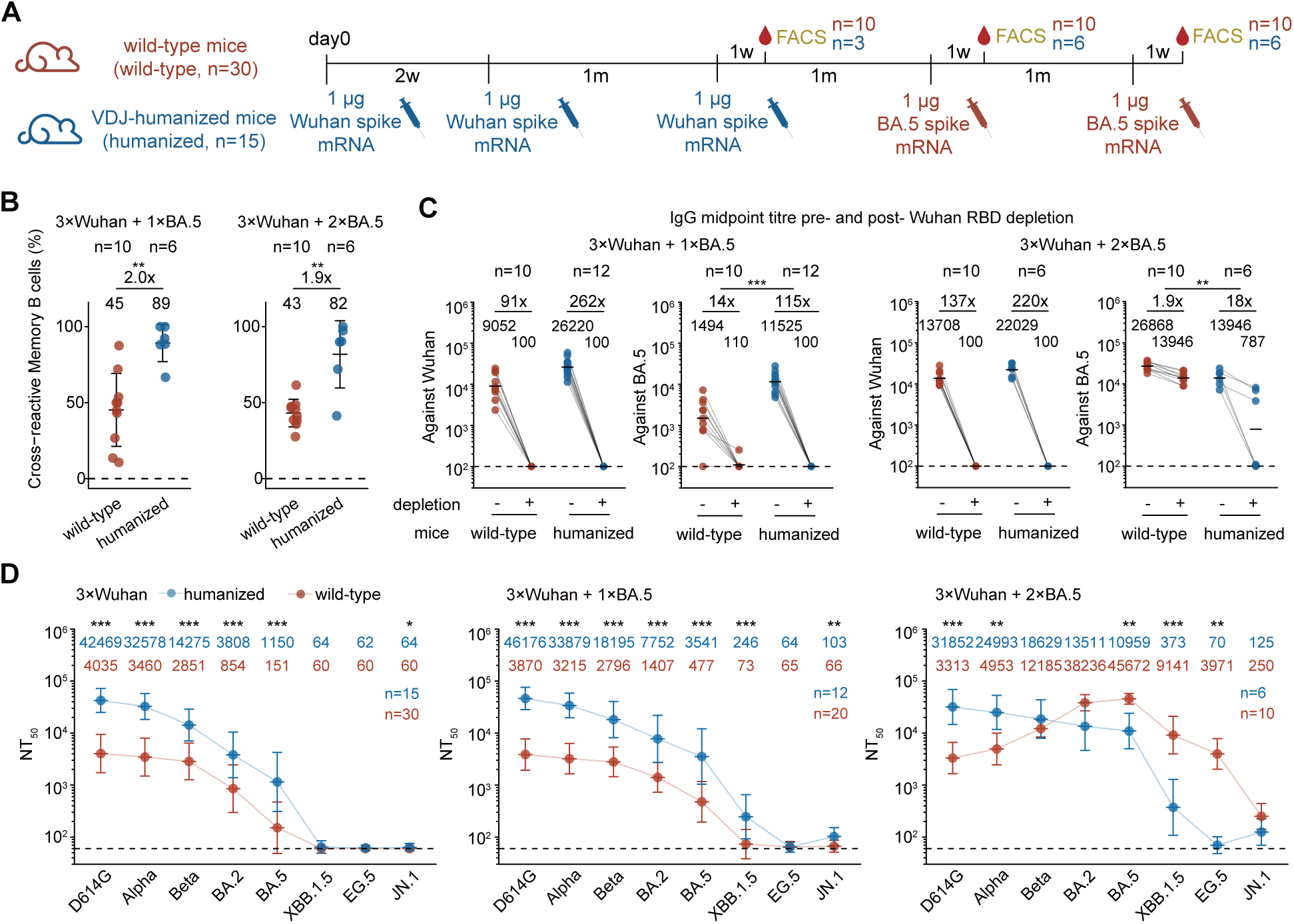
V(D)J-humanized mice recapitulate severe SARS-CoV-2 imprinting. (A) Schematic of the immunization regimen and sampling timeline for wild-type and V(D)J-humanized mice. The number of mice is indicated at the time point for each endpoint experiment. (B) Scatter plots showing the proportion of cross-reactive memory B cells in draining lymph nodes of the two mouse strains after one (left) or two (right) BA.5 boosts. Data are presented as mean ± standard deviation (SD). (C) Serum IgG midpoint titre of the two mouse strains after one (left) or two (right) BA.5 boosts against Wuhan or BA.5 RBD before and after Wuhan RBD depletion. Geometric mean values are displayed as bars and indicated above each group of data points. Statistical significance of the fold-reduction in titres was assessed between humanized and wild-type mice. Dashed lines indicate the limit of detection (midpoint titre = 100). (D) Serum neutralization titres (NT_50_) of the two mouse models after Wuhan priming (left), one BA.5 boost (middle), or two BA.5 boosts (right) against a panel of SARS-CoV-2 variant pseudoviruses. Geometric mean titres (GMTs) are shown on the top. Dashed lines indicate the limit of detection (NT_50_ = 60). Data are presented as geometric mean titres (GMT), with error bars indicating geometric standard deviation. Two-tailed Wilcoxon rank-sum tests were used in (B-D).

We first performed FACS analysis on mouse lymph nodes to assess cross-reactivity among memory B cells and germinal-center (GC) B cells (Supplementary Information Figure 1A). As expected, wild-type mice harbored abundant BA.5-specific memory and GC B cells after each BA.5 booster (Figures 1B and S1A). In striking contrast, the V(D)J-humanized mice exhibited a strong imprinting phenotype, developing a high proportion of cross-reactive memory B cells (89% and 82% after the first and second boosters, respectively), a phenotype consistent with severe imprinting described in strongly imprinted individuals. Despite similar overall percentages of GC B cells, humanized mice showed a significantly higher frequency of cross-reactive GC B cells after one BA.5 booster, indicating that their GC response was dominated by Wuhan-strain imprinted B cells (Figures S1A and S1B). Additionally, they presented a significantly higher frequency of class-switched memory B cells, which aligns with the preferential recall and expansion of a pre-existing memory population typical of immune imprinting (Figure S1C).

The imprinting was also faithfully reflected in the serological response. Wuhan RBD-depletion caused a substantially greater drop in BA.5-binding IgG titres in humanized mice than in wild-type mice (18-fold vs. 1.9-fold), although the magnitude of this effect varied among individual humanized mice (Figure 1C). Specifically, titres in three humanized mice dropped to undetectable levels, while the remaining three exhibited modest reductions. The post-depletion titre ratio was inversely correlated with the proportion of cross-reactive memory B cells (Supplementary Information Figure 3A), demonstrating consistency between cellular and serological assays and reflecting individual heterogeneity in imprinting strength.

We next compared serum neutralization profiles between the two mouse models (Figures 1D and S1D). Initially, following Wuhan priming, humanized mice mounted a significantly more potent and broad response than wild-type mice, consistent with the efficient induction of human germline-encoded public antibody responses. Two BA.5 boosters successfully shifted the neutralizing antibody (NAb) preferences of wild-type mice toward Omicron variants. In contrast, V(D)J-humanized mice retained a strongly ancestral-biased profile, with titres remaining highest against D614G, indicating strong immune imprinting that could not be overcome by two doses of BA.5 boosting (Figures 1D and S1D). Together, these data collectively demonstrate that humanized mice can mirror SARS-CoV-2 imprinting. This divergence between the two mouse models strongly suggests that the presence of human immunoglobulin V(D)J genes is a critical determinant of the imprinting phenotype.

### Distinct antibody landscapes of mouse models

To dissect the molecular basis of imprinting, we tracked the evolution of antibody repertoire from the two mouse models throughout Wuhan-priming and BA.5-boosting. We sorted RBD-specific memory B cells from spleens of humanized and wild-type mice following Wuhan priming (sorted on Wuhan RBD) and after each BA.5 booster (sorted on BA.5 RBD; Supplementary Information Figure 1B). Single-cell V(D)J sequencing of paired heavy- and light-chain variable regions generated 583 unique monoclonal antibodies (mAbs) from the six mice groups. These mAbs were recombinantly expressed as human IgG1 (Table S1), and their half-maximal inhibitory concentration (IC_50_) measured via pseudovirus neutralization assay (Figure S2A). Consistent with the established imprinting signature, enzyme-linked immunosorbent assay (ELISA) revealed that antibodies from humanized mice exhibited a higher proportion of cross-reactivity than those from wild-type mice (Figure S2B).

To systematically dissect the divergence in antibody epitope distribution between humanized and wild-type mice, we employed high-throughput yeast-display-based deep mutational scanning (DMS) to map the RBD mutations that could escape the isolated mAbs and define the epitope targeted ^14,23–26^. We built single-site saturation mutant libraries based on the Wuhan-Hu-1 and BA.5 RBDs and performed DMS for antibodies isolated following Wuhan priming or BA.5 boosting, respectively. The resulting escape profiles of 583 mAbs define the critical residues that mediate immune evasion and facilitating their precise categorization into distinct epitope clusters.

IC_50_-weighted escape profiles show that neutralizing antibodies from wild-type and humanized mice exhibit distinct escape hotspot sites (Figures 2A-2C and S2C-2E). Following Wuhan priming, the NAbs elicited in humanized mice substantially focused on Class 1 epitopes, featured by hotspots at residues 417, 456, 460, and 473. Although wild-type mice displayed a comparable landscape after Wuhan priming, they lacked these Class 1 peaks, instead featuring residue 477 as a characteristic hotspot (Figure 2A). BA.5 boosting leads to further divergence between the two mouse models. In humanized mice, the prominent Class 1 hotspots (e.g., residue 456) persisted obstinately throughout the boosting regimen. In contrast, the wild-type repertoire diverged significantly upon boosting, characterized by the emergence of distinct Class 1/4 escape peaks, such as residue 504 (Figures 2B and 2C).

**Figure 2.**
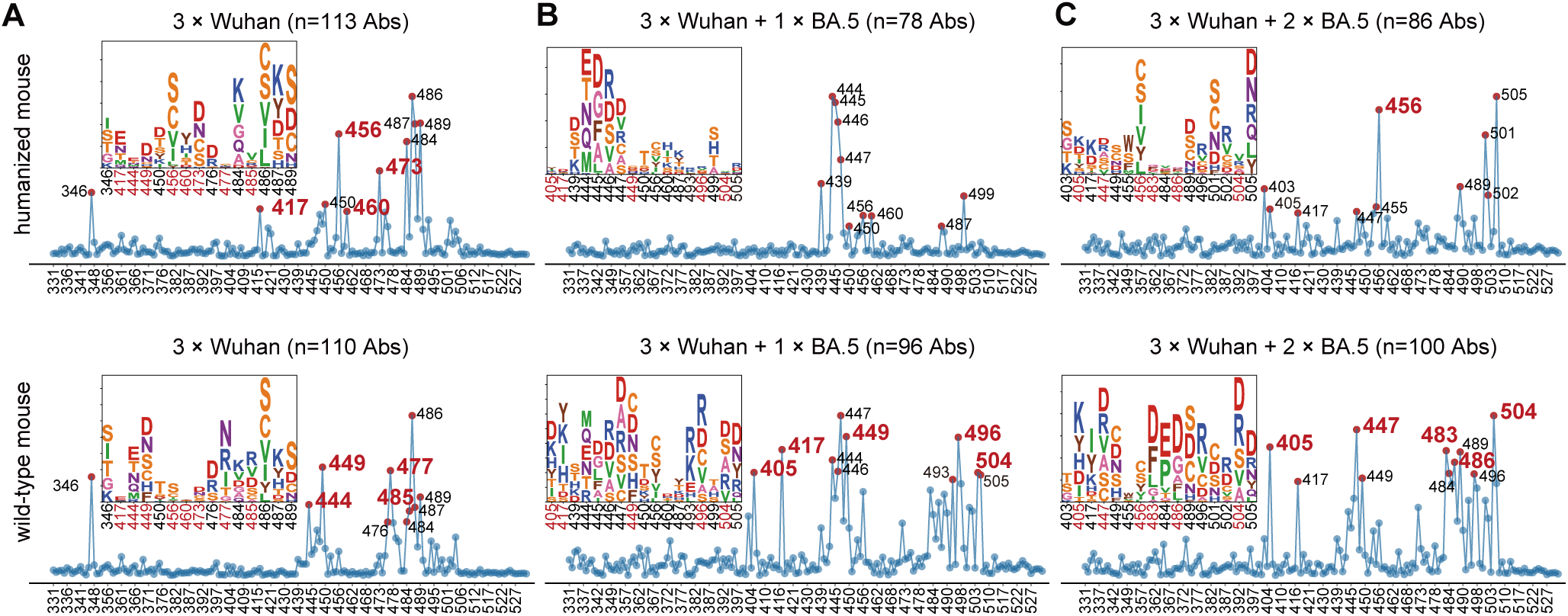
Distinct antibody landscapes between mRNA vaccinated humanized and wild-type mice. (A–C) Normalized average DMS escape scores for mAbs isolated from humanized (top) and wild-type (bottom) mice following Wuhan priming (A), the first BA.5 booster (B), or the second BA.5 booster (C). Escape scores were aggregated and weighted by the IC_50_ of each individual mAb (To focus on the neutralizing mAbs, lower IC_50_ corresponds to greater weight) against D614G for Wuhan-primed groups or BA.5 for BA.5-boosted groups. Codon constraints were applied (see Methods). In each panel, the ten residues with the highest cumulative escape scores are annotated, with their specific mutational escape profiles visualized as logo plots above the scatter plots. To highlight divergent immune pressure, residues exhibiting pronounced differences between humanized and wild-type strains (as identified in Figures S2C-S2E) are colored red in both the scatter plots and logo plots.

To translate these differences in escape sites into a more intuitive view of epitope-level repertoire differences, we categorized the 583 mAbs into 9 epitope groups (Supplementary Information Figure 2A) ^14^. Names of the epitope groups were generally assigned in line with the epitope groups on Wuhan RBD defined previously ^27,28^. We then comprehensively analyzed the antibody repertoires elicited in humanized and wild-type mice after each timepoint (Figures 3A, 3B and S3A-3D). Following Wuhan priming, humanized mice uniquely elicited a dominant population of antibodies targeting the A1 epitope (Figure 3A). Notably, antibodies targeting this epitope are closely related to the IGHV3-53 germline and target key residues such as 456, aligning with the escape hotspots. The foundational divergence between the two mouse models was established immediately after Wuhan priming. After two BA.5 boosting doses, while dynamic shifts occurred—including the abrogation of Group B and the emergence of Group A2—the A1 response remained persistent in humanized mice but was completely absent in wild-type mice (Figures 3C and S3A-S3E). In contrast, Omicron-specific antibodies in epitope groups B, D, and F3 contribute mostly to the neutralization against BA.5 exclusively in wild-type but not the V(D)J-humanized mice.

**Figure 3.**
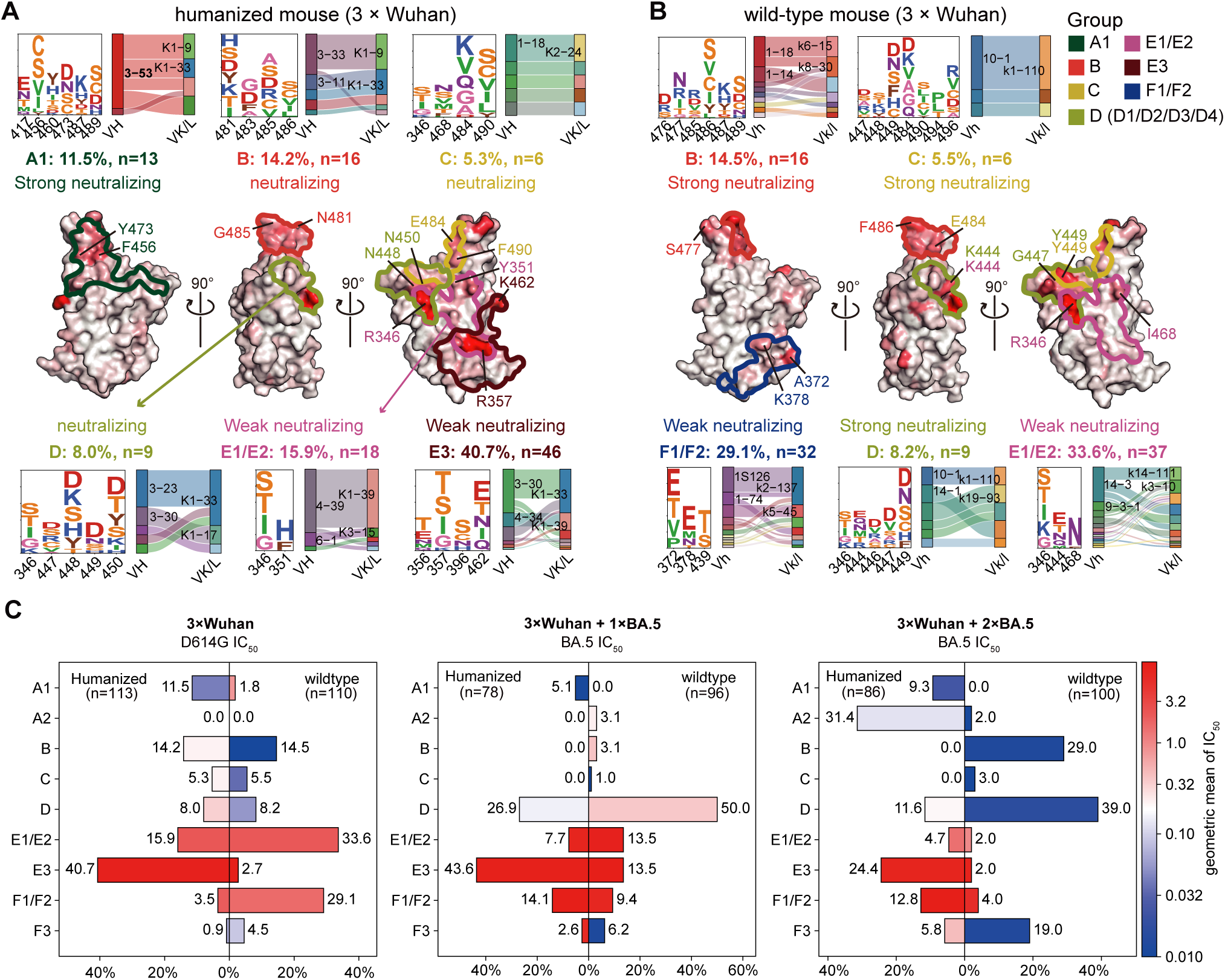
V(D)J germline difference shapes vaccination- induced SARS-CoV-2 antibody epitope distribution. (A and B) Epitope distribution of the antibody repertoire generated after Wuhan priming in humanized (A) and wild-type mice (B). The Wuhan RBD structure (PDB: 6m0j) is displayed as a surface map colored by normalized aggregate escape scores, with major epitope groups outlined in distinct colors. Their neutralizing category, antibody count, and percentage are indicated. Neutralizing category is classified based on geometric mean of IC_50_: Strong neutralizing (<0.1 µg/mL), Neutralizing (0.1 ≤ IC_50_ < 1 µg/mL), and Weak neutralizing (1 ≤ IC_50_ ≤ 10 µg/mL). Epitope groups accounting for <5% of the total antibodies are not labeled. Key escape residues for each group are displayed as logos, and the sites with the highest escape scores per group are labeled on the structure. Paired heavy and light chain V-gene usage for each epitope is shown in Sankey plots. (C) Pyramidal bar charts showing the proportional distribution of epitope groups in antibodies isolated from humanized and wild-type mice after Wuhan priming and after one or two BA.5 boosts. Bars are colored according to the log10 geometric mean IC_50_ of antibodies within each group.

Overall, DMS analysis demonstrates that Class 1 (A1) antibodies should play a pivotal role in the response to Wuhan priming, serving as a defining characteristic of the V(D)J-humanized mouse model, compared to the wild-type mice. This antibody subset substantially contributed to the polyclonal neutralization landscape and could be robustly recalled upon BA.5 boosting. Notably, these A1 antibodies exhibited remarkable germline convergence, consistently dominated by IGHV3-53 after both Wuhan priming and subsequent BA.5 boosting (Figure 3A). This is consistent with the enrichment of IGHV3-53/66-encoded Class 1 antibodies in strongly imprinted humans. Collectively, these findings led us to hypothesize that the dominant, IGHV3-53/66-driven A1 antibody response induced by Wuhan priming is the mechanistic driver of SARS-CoV-2 immune imprinting.

### Molecular mechanism of imprinting

Given that immune imprinting relies on the recall of pre-existing memory B cells, we then confirmed the cross-reactivity of these IGHV3-53/66-encoded A1 antibodies to determine their potential for reactivation by the Omicron booster (Figure 4A). 38% of A1 mAbs from Wuhan-primed humanized mice were cross-reactive, constituting a pre-existing memory pool that was efficiently recalled to achieve 100% cross-reactivity upon BA.5 boosting. In contrast, the negligible A1 response in primed wild-type mice was entirely Wuhan-specific and thus completely escaped by BA.5, resulting in the total absence of A1 antibodies following the boosters. Competitive SPR mapping revealed that A1 antibodies compete with ACE2 and the majority of antibodies targeting neutralizing epitopes (A2, B, C, D4, F3). Conversely, they showed minimal to no competition with Group D1 or the weakly or non-neutralizing E and F antibodies (Figure 4B).

These results indicate that A1 antibodies can sterically mask most other major neutralizing sites on Omicron RBD from access by other antibodies and their corresponding B cell receptors. Enabled by the IGHV3-53/66 germline genes, which encode a prototypic public Class 1 antibody response targeting the ACE2-binding site ^29–36^, Wuhan priming induces a high-frequency pool of these potent “masking” antibodies, the abundance of which scales directly with the intensity of the priming. Critically, a significant portion of this pool is not escaped by Omicron and remains cross-reactive, making it available for recall upon subsequent Omicron exposure. Therefore, during Omicron vaccination, this pre-existing and cross-reactive memory B cell population is preferentially reactivated and expanded. Such a potent, recalled response could actively outcompete and suppress the *de novo* activation of B cells targeting novel Omicron-specific epitopes by antibody masking, providing a direct mechanistic basis for the strong immune imprinting observed in humanized mice (Figure 4C).

To test whether these antibodies can indeed drive strong imprinting *in vivo*, we performed passive antibody transfer experiments in wild-type mice (Figure 4D). Wuhan-primed wild-type mice were infused with BD55-1205 (a representative IGHV3-66-encoded A1 broad neutralizing antibody ^37^) at various doses and formats (human or mouse IgG1) one day prior to BA.5 boosting (Figures 4D). Controls received either PBS or a non-neutralizing antibody (BD57-2665, a representative F1 antibody targeting the cryptic sites of RBD, Figure 4E). Flow cytometry analysis of lymph nodes showed that mice receiving 400 µg or 200 µg BD55-1205 hIgG1 and 200 µg BD55-1205 mIgG1 developed pronounced imprinting, evidenced by significantly elevated frequencies of cross-reactive memory B cells at both post-boost time points compared with PBS controls (Figure 4F). After the first BA.5 boost, GC B cells in BD55-1205–treated mice also showed markedly higher cross-reactivity, demonstrating that early maturation and clonal expansion within the GC was dominated by recalled, imprint-driven B cells (Figure 4G). In contrast, BD57-2665 mIgG1–treated mice showed no significant change in cross-reactive memory B cell or GC B cell frequencies, indicating that non-neutralizing antibody blockade does not induce imprinting, ruling out the possibility that the transferred antibody simply depleted the vaccine antigen and prevented a successful BA.5 immunization (Figures 4F and 4G). Correspondingly, Wuhan-RBD depletion of serum from BD55-1205–treated mice produced a more severe drop in BA.5 IgG titres, whereas BD57-2665 mIgG1 and PBS groups showed minimal titre reduction (Figure 4H). Notably, the mIgG1 format induced a significantly stronger imprinting phenotype than hIgG1, which is likely attributable to the species-matched Fc region (Figures 4F-4H). Pharmacokinetic analysis confirmed that the passively transferred antibodies had decayed to near-undetectable levels by the time of analysis, thus eliminating their potential interference with ELISA measurements (Figures S4A and S4B). Importantly, we demonstrated that the suppressive effect of BD55-1205 on the BA.5-specific response was dose-dependent, with doses as low as 12.5 µg being sufficient to achieve inhibition (Figures S4C and S4D). Together, these results confirm that IGHV3-53/66-encoded A1 neutralizing antibodies can suppress the emergence of Omicron-specific B cell and antibody responses to induce severe imprinting *in vivo*.

This explains why immune imprinting is recapitulated in V(D)J-humanized mice but not in wild-type mice—only the former harbor this human germline gene required to produce such imprinting-prone antibodies. Similarly, consistent with our observation in human cohorts ^11^, in mRNA-vaccinated individuals, the potent Wuhan priming response by mRNA vaccination (elicited by monovalent Wuhan vaccines and, in certain populations, bivalent Wuhan/BA.1 or Wuhan/BA.5 vaccines) could induce a larger pool of IGHV3-53/66-encoded A1 antibodies, thereby establishing pronounced imprinting. In contrast, recipients of inactivated vaccines typically experience weaker priming responses and, following stringent containment policies during 2021-2022, undergo substantial waning of Wuhan antibodies. This attenuated pre-existing immunity, potentially leading to lower IGHV3-53/66 antibody abundance, fails to fully suppress subsequent Omicron-specific responses, resulting in more flexible and adaptive humoral evolution.

### IGHV3-53 knock-in mouse model

Since wild-type mice could not recapitulate human SARS-CoV-2 immune imprinting, evaluating SARS-CoV-2 vaccine updates and broad-spectrum vaccine design in mice is highly problematic. Although V(D)J-humanized mice could capture human imprinting phenotypes, its significant cost creates a pressing need for a more accessible and cost-effective model for vaccine evaluation. Building on our finding that the human IGHV3-53/66 germline is the primary driver of immune imprinting, we engineered a knock-in mouse model by replacing the murine Ighv3-1 gene with human IGHV3-53 (Supplementary Information Figure 4B). This generated both heterozygous (IGHV3-53^+/-^) and homozygous (IGHV3-53^+/+^) mice. High-throughput Genome-wide Translocation Sequencing (HTGTS) of the naïve B cell repertoire confirmed the successful knock-in, revealing IGHV3-53 usage at 0.5% in heterozygous and 1.5% in homozygous mice, with a corresponding ablation of Ighv3-1 usage (0.1% and 0.0%, respectively, Figure 5A).

**Figure 4.**
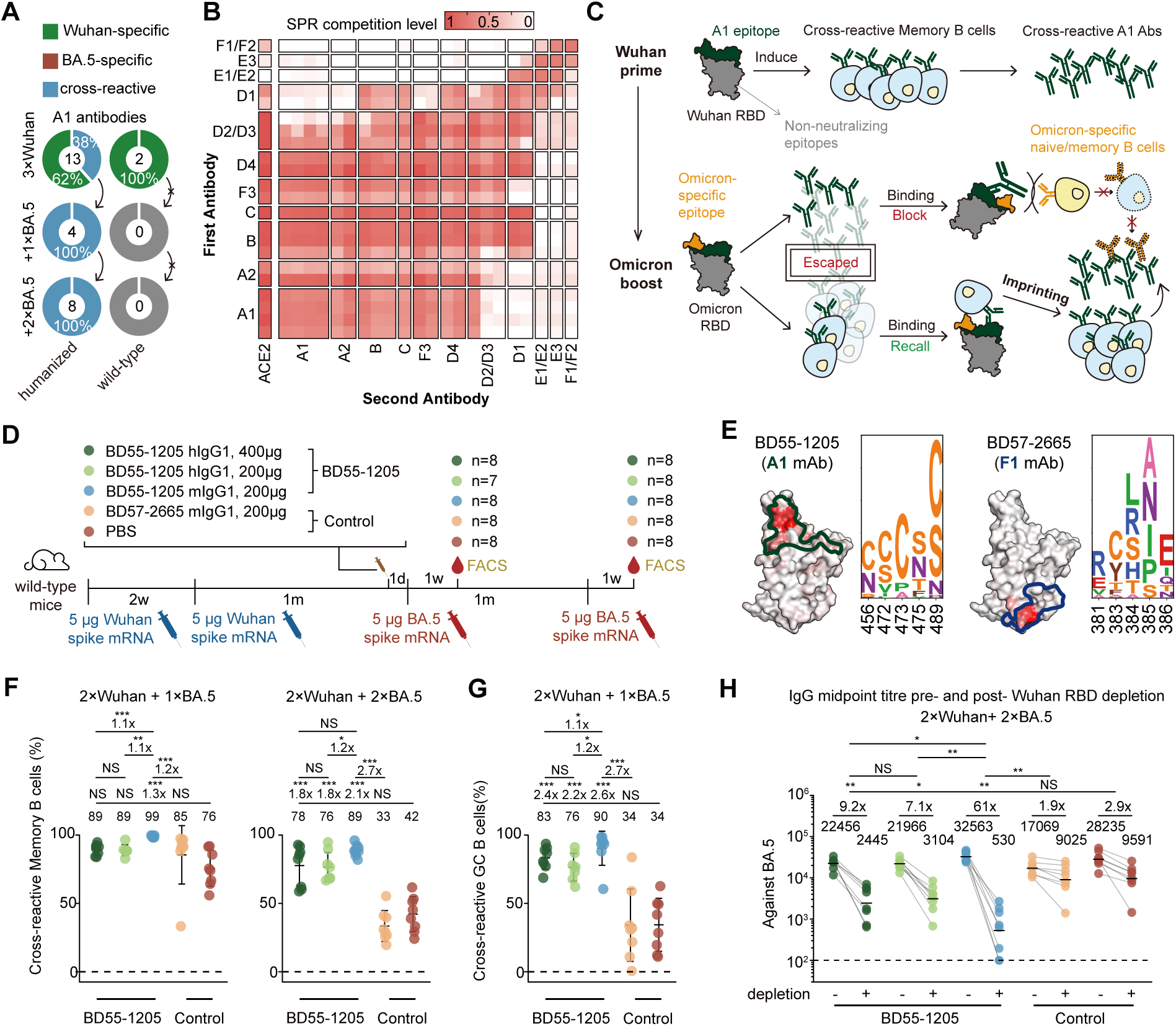
IGHV3-53/66-encoded Class 1 antibody drives SARS-CoV-2 imprinting through epitope masking. (A) Donut plots showing the cross-reactivity of A1 antibodies from humanized or wild-type mice. The number of antibodies is indicated in the centre of the donut. Antibodies exhibiting ELISA OD450 values > 2 against both WT and BA.5 RBDs (1 μg/mL) were defined as cross-reactive. Those showing an OD450 > 2 for one variant but < 2 for the other were classified as specific. (B) Heatmap of competitive SPR for various antibody groups. The definition of the competition score is described in the Methods section. (C) Schematic of the molecular mechanism by which pre-existing IGHV3-53/66-encoded A1 antibodies cause strong immune imprinting. (D) Schematic of the antibody passive transfer experiment. Timing of mRNA vaccinations, antibody injection, blood collection, and FACS analysis are indicated. Mice were divided into experimental groups (receiving 400 µg BD55-1205 hIgG1, 200 µg BD55-1205 hIgG1, or 200 µg BD55-1205 mIgG1) and control groups (receiving 200 µg BD57-2665 mIgG1 or PBS). The number of mice per group is indicated at the endpoint. (E) DMS escape map logoplots for BD55-1205 and BD57-2665 and their projection onto the SARS-CoV-2 Wuhan RBD (PDB: 6m0j). (F) Scatter plots showing the proportion of cross-reactive memory B cells in draining lymph nodes after one (left) or two (right) BA.5 boosts. (G) Scatter plots showing the proportion of cross-reactive germinal center B cells in draining lymph nodes after one BA.5 boost. Data are presented as mean ± standard deviation (SD). (H) Serum IgG midpoint titre against BA.5 RBD before and after Wuhan RBD depletion. Geometric mean values are displayed as bars and indicated above each group of data points. Statistical significance of the fold-reduction in titres was assessed between groups. Dashed lines indicate the limit of detection (midpoint titre = 100). Two-tailed Wilcoxon rank-sum tests were used in (F-H).

**Figure 5.**
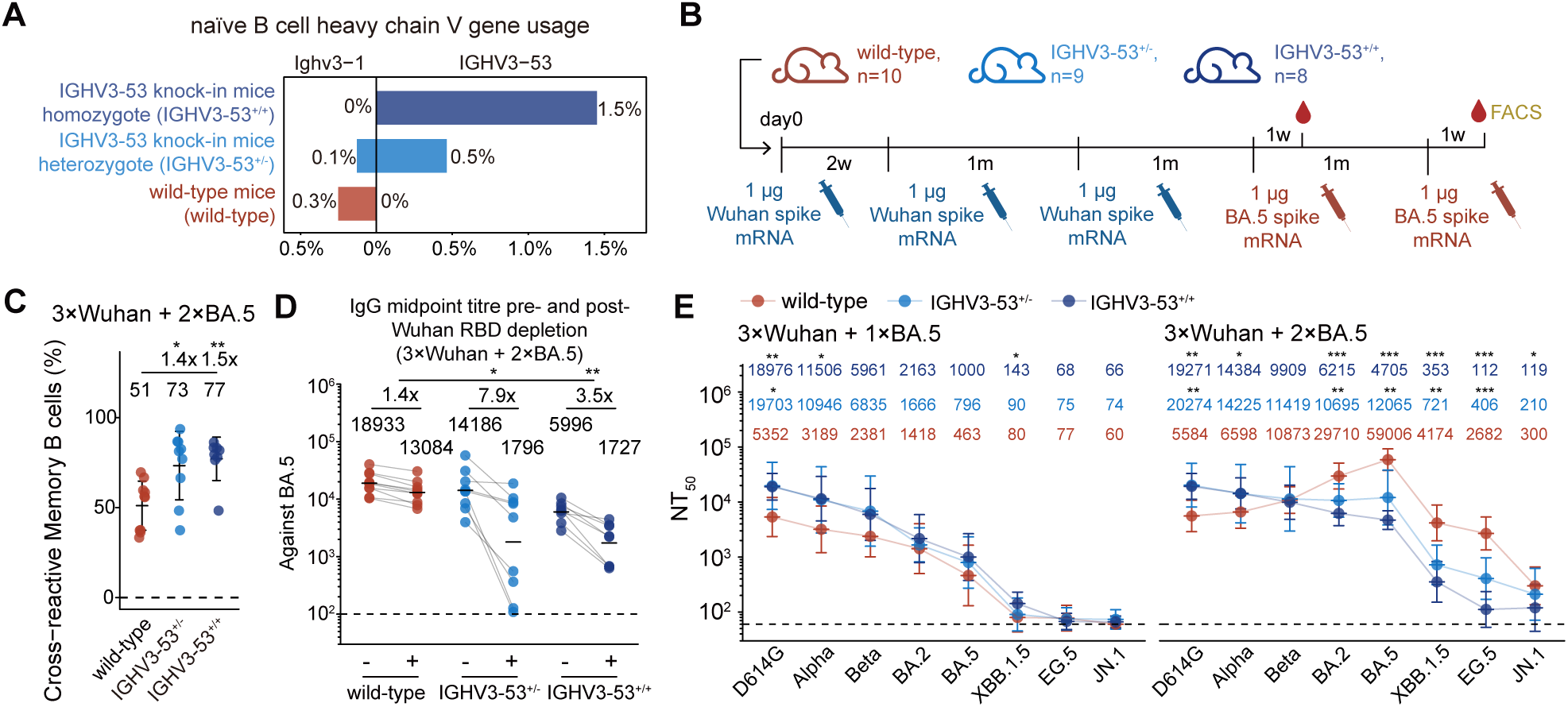
IGHV3-53 knock-in was sufficient to induce severe imprinting in wild-type mice. (A) Usage percentage of IGHV3-53 and Ighv3-1 in naïve B cells from HTGTS sequencing of IGHV3-53^+/+^ (n=3), IGHV3-53^+/-^ (n=3), and wild-type (n=3) mice. Genomic DNA from mice within each group was pooled for sequencing. (B) Schematic of the immunization regimen and time points for blood collection and FACS analysis of the three mouse models involved in this study. The number of mice is indicated above the timeline. (C) Scatter plots showing the proportion of cross-reactive memory B cells in draining lymph nodes of the three mouse models after two BA.5 boosts. Data are presented as mean ± standard deviation (SD). (D) Serum IgG midpoint titre of the three mouse strains after two BA.5 boosts against BA.5 RBD before and after Wuhan RBD depletion. Geometric mean values are displayed as bars and indicated above each group of data points. Statistical significance of the fold-reduction in titres was assessed between groups. Dashed lines indicate the limit of detection (midpoint titre = 100). (E) Serum neutralization titres (NT_50_) of the three mouse strains after one (left) or two (right) BA.5 boosts against a panel of SARS-CoV-2 variant pseudoviruses. Geometric mean titres (GMTs) are shown on the top. Dashed lines indicate the limit of detection (NT_50_ = 60). Data are presented as geometric mean titres (GMT), with error bars indicating geometric standard deviation. Two-tailed Wilcoxon rank-sum tests were used in (C-E).

To test the model, we immunized wild-type, heterozygous, and homozygous mice using the same regimen of three Wuhan primes followed by two BA.5 boosts (Figure 5B). Subsequent flow cytometry analysis and depletion assays confirmed that both IGHV3-53^+/-^ and IGHV3-53^+/+^ mice faithfully recapitulated the strong immune imprinting phenotype (Figures 5C and 5D). The serum neutralization profiles of IGHV3-53^+/-^ and IGHV3-53^+/+^ mice also showed a trend remarkably similar to that of the V(D)J-humanized mice (Figures 5E and S5A). These results demonstrate that both heterozygous and homozygous IGHV3-53 knock-in mice faithfully recapitulate the strong immune imprinting phenotype.

We subsequently utilized this model to simulate the real-world SARS-CoV-2 exposure history characteristic of mRNA-vaccinated populations (Figure 6A). Specifically, mice received a two-dose Wuhan priming series, mimicking the standard primary course. This was followed by sequential boosters with bivalent Wuhan/BA.1 and Wuhan/BA.5 vaccines, a monovalent XBB.1.5 vaccine, and a monovalent JN.1 vaccine, corresponding to the recommended mRNA booster updates during the Omicron era. These regimens effectively reconstruct the antigenic trajectory of SARS-CoV-2 evolution encountered by humans. Compared to wild-type controls, IGHV3-53^+/+^ mice recapitulated a more pronounced immune imprinting phenotype following the six-dose regimen. Specifically, while retaining potent neutralization against D614G and early Omicron variants (antecedent to JN.1, Figure 6B), IGHV3-53^+/+^ mice exhibited a compromised breadth of neutralization against the newly emerged JN.1 sublineages. Specifically, titres against KP.2, KP.3, LP.8.1.1, NB.1.8.1, and XFG were significantly suppressed compared to those in wild-type mice, indicating an inhibition of *de novo* responses to the newly emerged Omicron-specific epitopes in these variants. Notably, IGHV3-53^+/+^ mice showed higher neutralization titres against BA.3.2.2 than wild-type mice (Figure 6B), consistent with the known sensitivity of BA.3.2.2 to IGHV3-53/66-encoded Class 1 antibodies ^11^. Similar trends were observed after five doses (Figure S6A). Furthermore, comparing neutralization profiles pre- and post-JN.1 boost revealed a strong back-boosting effect in IGHV3-53^+/+^ mice, where JN.1 immunization significantly elevated titres against antecedent strains (pre-JN.1). In contrast, wild-type mice displayed no such effect and even exhibited a significant decline in D614G titres (Figure S6B).

**Figure 6.**
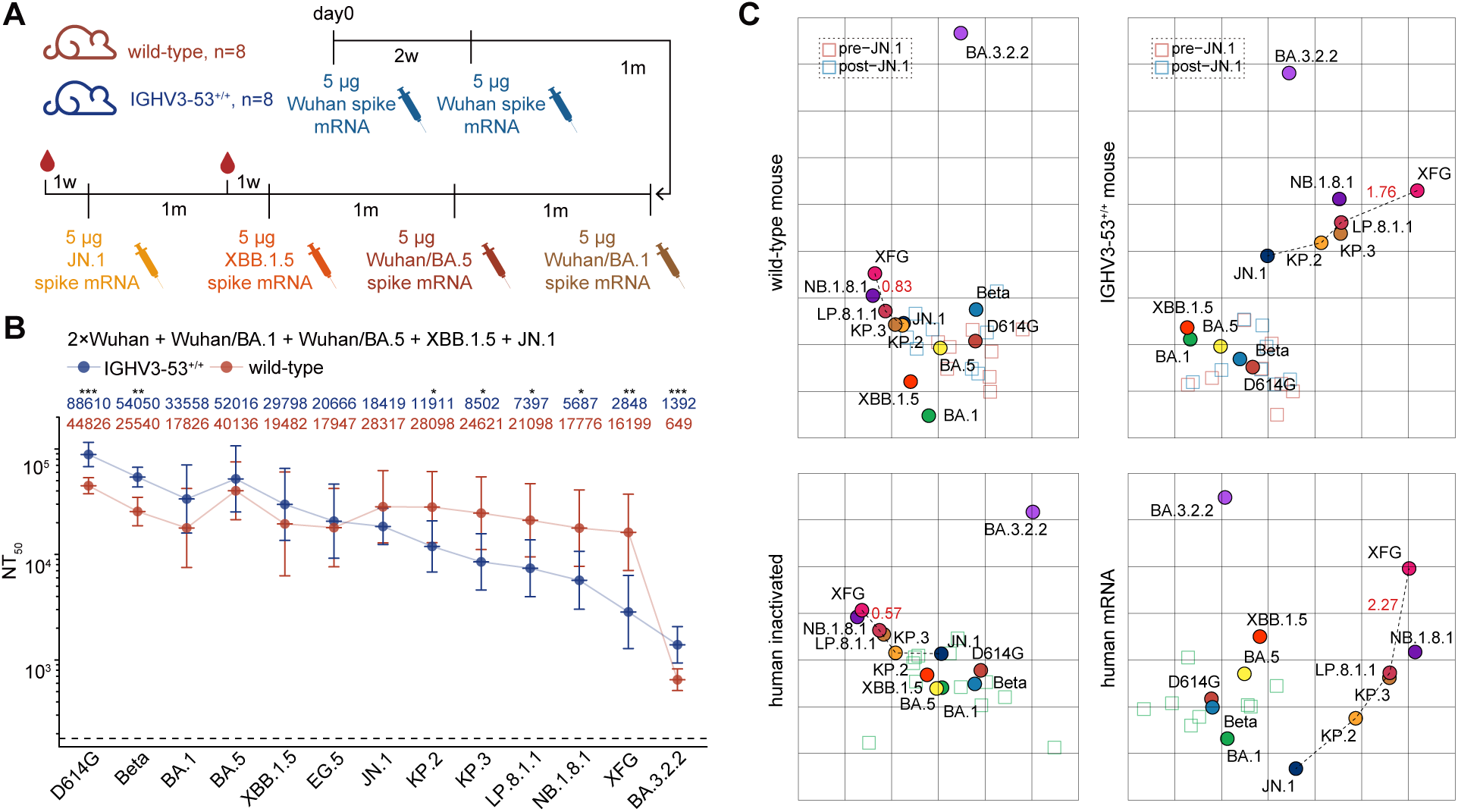
IGHV3-53 knock-in mice faithfully reflect SARS-CoV-2 antibody map in human. (A) Schematic of the immunization regimen simulating real-world SARS-CoV-2 exposure history. (B) Serum neutralization titres (NT_50_) of IGHV3-53^+/+^ and wild-type mice against a panel of SARS-CoV-2 variant pseudoviruses. Geometric mean titres (GMTs) are shown on the top. Dashed lines indicate the limit of detection (NT_50_ = 180). Data are presented as geometric mean titres (GMT), with error bars indicating geometric standard deviation. (C) Antigenic cartography was performed using mouse (top) and human (bottom) serum neutralization data. Each square indicates a serum sample, and each circle indicates a SARS-CoV-2 variant. Human data were obtained from previous study ^11^. Two-tailed Wilcoxon rank-sum tests were used in (B).

Consequently, the two mouse models exhibited distinct susceptibility to viral escape. Following XBB.1.5 immunization, neutralization against EG.5 was significantly reduced compared to XBB.1.5 in IGHV3-53^+/+^ mice, a reduction not seen in wild-type mice (Figure S6C). Similarly, after the JN.1 boost, IGHV3-53^+/+^ mice showed significant susceptibility to escape by KP.2 and KP.3 relative to JN.1, whereas wild-type mice did not. Consistently, the fold-reduction in titres against XFG relative to NB.1.8.1 was greater in IGHV3-53^+/+^ mice than in wild-type mice (Figure S6C). These results demonstrate that an IGHV3-53-driven imprinted response intrinsically creates predictable vulnerabilities to escape by specific Omicron sublineages such as XFG.

To visualize how IGHV3-53^+/+^ mice could help with SARS-CoV-2 vaccine update evaluation, we constructed antigenic cartography using serum neutralization data from both mouse models and reference human cohorts (Figure 6C) ^11^. The resulting maps revealed striking topological differences. Wild-type mice exhibited a ’condensed’ antigenic landscape similar to the inactivated-only cohort, characterized by short antigenic distances between all JN.1 sublineages. In contrast, the antigenic map of IGHV3-53^+/+^ mice closely mirrors that of mRNA-vaccinated cohort, with both displaying substantial antigenic distances between JN.1, KP.2, KP.3/LP.8.1.1, and XFG, supporting vaccine antigen updates. Crucially, only the IGHV3-53^+/+^ model faithfully reproduces the substantial antigenic distance between XFG and LP.8.1.1 characteristic of the mRNA-vaccinated cohort. These findings further implicate the IGHV3-53 knock-in mice could faithfully recapitulates human SARS-CoV-2 immune imprinting, and can serve as a superior model for preclinical SARS-CoV-2 vaccine evaluation.

## Discussion

Overall, our study establishes that the abundance of pre-existing IGHV3-53/66-encoded Class 1 antibodies is the primary determinant of SARS-CoV-2 immune imprinting severity. When ancestral priming establishes an abundant pool of cross-reactive IGHV3-53/66-encoded A1 antibodies, subsequent Omicron exposure preferentially recalls this response and suppresses de novo Omicron-specific antibodies through epitope masking. Conversely, when this antibody route is absent or insufficient, Omicron boosting can more readily elicit adaptable variant-focused responses.

We acknowledge the individual heterogeneity within our humanized mouse model, where a part of them exhibited weaker immune imprinting. We hypothesize that the isolated Omicron-specific A2 antibodies likely derived from these mice. Although the pooling of samples during B cell sorting prevents us from retrospectively tracing specific antibodies to individual animals, the existence of these outlier responses does not undermine our core conclusions that humanized mice collectively develop a significantly stronger imprinting phenotype in both serum and memory B cell compartments. These “outlier” mice likely resemble inactivated vaccine recipients, where the abrupt exposure to BA.5 resulted in extensive escape of pre-existing A1 antibodies, leaving insufficient cross-reactive binding to effectively mask Omicron-specific epitopes. In contrast, sequential exposure to BA.1/BA.2 in mRNA vaccinees involved a gradual antigenic shift, allowing A1 antibodies to mature and expand without facing the drastic escape seen with BA.5. This preserved a sufficient cross-reactive pool to effectively mask the *de novo* response upon the subsequent BA.5 exposure.

Importantly, we developed an accessible and robust IGHV3-53 knock-in mouse model that overcomes the limitations of wild-type mice, which fail to recapitulate the SARS-CoV-2 imprinting phenotype. By simulating real-world SARS-CoV-2 exposure histories in this model, we successfully mapped the humoral response landscape of mRNA human vaccinees in mice. Moving forward, this model should be leveraged to conduct parallel evaluations of boosting regimens derived from diverse, newly emerging variants. Such comparative studies will provide the essential data support required to guide the further updating and optimization of SARS-CoV-2 vaccines. Furthermore, as a platform that explicitly exposes the ’imprinted barrier’ of mRNA recipients, it would enable the future research of vaccine candidates capable of overcoming such entrenched imprinting.

## Acknowledgments

This project is financially supported by Changping Laboratory (2026D-04-01 to Y.C.).

## Data and code availability

Information of isolated mAbs involved in this study have been included in the Supplementary Tables. All data presented in this manuscript are available from the lead contact, (Y.C.), upon a reasonable request under a completed Material Transfer Agreement. No custom code was developed for this study. Data analysis was performed using standard software or packages/libraries in R or python, as detailed in the Methods section.

## Author contributions

Y.C. designed and supervised the study. X.N., F.J. and Y.C. wrote the manuscript with input from all authors. X.N., W.S., R.A. and Y.W. performed B cell sorting, single-cell V(D)J sequencing experiments and data analysis. X.N., Y.L., K.L., S.L., R.K., X.C., R.A., and Y.W. performed FACS analysis. H.S. and F.J. obtained and analyzed the DMS data. J.W. and F.S. performed antibody expression. W.W. constructed mRNA vaccines and conducted mouse immunization. Y.Y. constructed the pseudotyped virus. L.Y. performed the pseudovirus neutralization assays, ELISAs and SPR experiments. Y.H., S.L., X.N. and Y.L. performed the HTGTS-Rep-seq and data analysis.

## Methods

### Mice

The animal experiments were conducted under protocols approved by Laboratory Animal Center of Peking University (Approval No. BIOPIC-CaoYL-1). The humanized mouse model (NeoMab-IgG), provided by NeoMab Biotechnology Co., Ltd., was created by in situ replacing the mouse heavy chain variable region genes and kappa light chain variable region genes with human genes in BALB/c mouse background. The mouse constant region genes were retained, making sure the Fc of the immunoglobulins can interact with the Fc receptors expressed on other immune cells normally, supporting standard immune system development and response. The IGHV3-53^+/+^ or IGHV3-53^+/-^mice were generated by Cyagen Biosciences (Suzhou, China). Mice of all strains (6–8 weeks old) were housed under specific-pathogen-free (SPF) conditions, maintained at 22 ± 2°C with 50–60% humidity on a 12-hour light/dark cycle. All animals had *ad libitum* access to standard chow and sterile water. Female mice were used for the VDJ-humanized and wild-type BALB/c models, while both sexes were used for the IGHV3-53 knock-in models.

### Cell lines

The Huh-7 (JCRB, 0403) and HEK293T (ATCC, CRL-3216) cell lines were cultured in DMEM (Hyclone) supplemented with 10% fetal bovine serum (FBS, Hyclone) and 1% penicillin-streptomycin at 37°C with 5% CO_2_. The Expi293F (ThermoFisher, A14527) cell line was cultured in 293F Hi-exp medium (OPM Biosciences) at 37°C with 5% CO_2_ and 175 rpm shaking.

### Generation of IGHV3-53 knock-in mice

To generate IGHV3-53 knock-in mice, the targeting construct replaced the sequence from ATG start codon to exon 3 of mouse Ighv3-1 by the sequence from ATG start codon to exon 2 of the human IGHV3-53, including the introns. The Cas9 protein, construct and guide RNAs were microinjected into zygotes from BALB/cAnCya wild-type mice. The embryos were transferred to recipient female mice to obtain F0 mice. The genotype of IGHV3-53^+/+^ or IGHV3-53^+/-^ mice was confirmed by PCR and sequencing.

### mRNA-LNP synthesis and formulation

A PSP73 plasmid bearing the antigen insert followed by a 120-nt poly(T) tract was linearized with the appropriate restriction enzyme. This DNA served as a template for *in vitro* transcription process to generate RNA that encoded the SARS-CoV-2 Wuhan, BA.1, BA.5, XBB.1.5 and JN.1 S6P (F817P, A892P, A899P, A942P, K986P, V987P, R683A and R685A) protein. Linearized DNA template (1 µg) was transcribed for 2 hours at 37°C using the EasyCap T7 Co-transcription Kit with CAG Trimer (Vazyme). Following transcription, the DNA template was digested by incubation with 1 U of RNase-free DNase I for 15 minutes at 37°C. The resulting mRNA was purified with VAHTS RNA Clean Beads (Vazyme). The concentration and purity of the purified mRNA were determined by UV spectrophotometry (absorbance at 260 nm and A260/A280), and its integrity was verified by agarose gel electrophoresis.

The mRNA was encapsulated in a functionalized lipid nanoparticle (LNP) as described previously^50^. The lipid mixture (oil phase) was prepared by dissolving SM-102, DSPC, cholesterol, and DMG-PEG2000 in 100 % ethanol at a molar ratio of 50:10:38.5:1.5 to a total concentration of 9.04 mg/mL. The filter-sterilized (0.22 µm) aqueous phase, consisting of 50 mM sodium citrate buffer (pH 4.0), and the purified mRNA was diluted within it to a concentration of 133 µg/mL. Oil and aqueous phases were then rapidly mixed at a 3:1 volume ratio using a staggered herringbone micromixer, operating at a total flow rate of 12 mL/min to induce LNP self-assembly. The crude LNP suspension underwent buffer exchange and purification via overnight dialysis at 4°C against a solution of 10 mM Tris-HCl (pH 7.4) with 8-10% (w/v) sucrose, using 10 kDa MWCO dialysis cassettes. The dialyzed LNPs were subsequently concentrated by centrifugation at 1,500-2,500 × g for 10-15 minutes at 4°C using 10 kDa MWCO ultrafiltration units (ULRC0100150P). The final formulation was adjusted to a sucrose concentration of 8.7% (w/v), aliquoted, flash-frozen in liquid nitrogen, and stored at -80°C.

The final product was subjected to rigorous quality control. Key attributes including particle size (by dynamic light scattering), RNA encapsulation efficiency (by RiboGreen assay), mRNA integrity (by capillary electrophoresis), osmolality, and endotoxin levels (by LAL assay) were assessed. All manufactured batches were required to meet the following release criteria: a particle diameter of 80-100 nm, ≥90% RNA encapsulation, ≥80% mRNA integrity, and an endotoxin level below 1 EU/mL.

### Mouse immunization

For immunization, mice were administered mRNA-LNP vaccines. The specific immunization regimens, including vaccine type, dosage, and timelines, are detailed in the schematics of Figures 1A, 4D, 5B, 6A, S4A and S4C. On the day of administration, LNP-mRNA vials were thawed on ice and diluted in sterile 1× PBS to the appropriate concentration. Each mouse was injected intramuscularly (i.m.) into the quadriceps muscle with 100 µL of the vaccine solution using a 29 G insulin syringe. To prevent leakage of the inoculum, the needle was held in place for 3–5 seconds post-injection. Due to the limited availability of the IGHV3-53 knock-in mice, the sample size and sex distribution varied slightly across the experimental groups shown in Figure 6. Sex distribution was 4 males and 4 females for IGHV3-53+/+ mice, 3 males and 6 females for IGHV3-53+/- mice in figure 5B, and 4 males and 4 females for IGHV3-53+/+ mice in figure 6A.

### Passive antibody transfer

For passive antibody transfer, purified mAbs (BD55-1205 mIgG1, BD55-1205 hIgG1, or BD57-2665 mIgG1) with low endotoxin levels (<0.02 EU/mg) were used. The antibodies were diluted in sterile PBS to the desired concentration for injection. Female BALB/c mice (6–8 weeks old) received a 200 µL dose via intraperitoneal (i.p.) injection in the left lower quadrant using a 26 G needle. Mice were monitored for at least 5 minutes following the injection, and any subsequent doses were administered according to the specified experimental schedules.

### Pseudovirus preparation and neutralization assay

We generated SARS-CoV-2 variant spike protein pseudovirus as described previously ^14,18,23,51–53^. Plasmids encoding a codon-optimized SARS-CoV-2 Spike (S) protein were constructed by inserting the S gene into the pcDNA3.1 vector. To produce pseudovirus, 293T cells were transfected with the S protein expressing plasmids with Lipofectamine 3000 (Invitrogen) and subsequently infected with G*ΔG-VSV (Kerafast). After 24 hours, the supernatant containing the pseudovirus was harvested, filtered through a 0.45 μm filter, aliquoted, and stored at -80°C.

Neutralization assays were performed using the Huh-7 cell line. Monoclonal antibodies or serum samples were serially diluted in DMEM and incubated with the pseudovirus in 96-well plates for 1 hour at 37°C with 5% CO₂. Following incubation, Huh-7 cells were seeded into the wells (2×10^4^ cells per well) and cultured for an additional 24 hours at 37°C with 5% CO₂. To assess infection levels, the culture supernatant was removed and left 100 μl in each well. The Bright-Lite Luciferase Assay Substrate was reconstituted with its corresponding Assay Buffer (Vazyme), and this mixture was added to the wells. After incubating in the dark for 2 minutes, luminescence was measured using a microplate spectrophotometer (PerkinElmer). The NT_50_ or IC_50_ values were determined using a three-parameter logistic regression model.

### Mouse tissue processing and B cell extraction

Following euthanasia, the spleen, inguinal lymph nodes, and popliteal lymph nodes were harvested and placed in RPMI 1640 culture medium (Invitrogen) containing 5% (v/v) FBS. Single-cell suspensions were prepared by mechanical disruption using the plunger of a syringe. The popliteal and inguinal lymph nodes from each mouse were pooled, ground, and filtered through a 40 µm cell strainer. For the spleen, tissue was processed by the same grinding method and passed through a 70 µm cell strainer, followed by centrifugation and lysis of red blood cells using 1× RBC Lysis Buffer (Invitrogen eBioscience). After washing steps and centrifugation, the resulting cell pellets were resuspended in PBS containing 2% (v/v) FBS.

Splenic B cells were enriched from the splenic single-cell suspensions via immunomagnetic negative selection with the EasySep™ Mouse Pan-B Cell Isolation Kit (STEMCELL). Following the manufacturer’s protocol, the untouched, purified B cells were collected and washed in PBS with 2% (v/v) FBS. The cell numbers of total lymph node cells and purified splenic B cells were determined using 0.4% (w/v) trypan blue stain and a Countess Automated Cell Counter.

### Flow cytometry analysis and antigen-specific B cell sorting

For the characterization of B cell responses in immunized mice, single-cell suspensions from the inguinal and popliteal lymph nodes were stained for flow cytometry analysis. The cells were stained with a panel including PE/Cyanine7 anti-mouse CD38, Brilliant Violet 605™ anti-mouse/human B220, APC/Cyanine7 anti-mouse IgD, Brilliant Violet 711™ anti-mouse IgM, and FITC anti-mouse/human GL7 (BioLegend). The antigen probe cocktail consisted of biotinylated BA.5 RBD conjugated with PE- and APC-streptavidin, and Wuhan RBD conjugated with BV421-streptavidin. Data for lymph node analysis were acquired on a Symphony A5SE cytometer (BD Biosciences).

For the isolation of antigen-specific mouse splenic B cells, purified splenic B cells were stained using the identical panel of antibodies and RBD probes as described for the lymph node analysis. MoFlo Astrios EQ Cell Sorter (Beckman Coulter) was used for all sorting experiments, targeting live (7-AAD⁻), B220⁺, CD38⁺, class-switched (IgM⁻ and IgD⁻), non-germinal center (GL7⁻) B cells that bound to the Wuhan or BA.5 RBD.

For all procedures, data were collected via Summit 6.0 software (Beckman Coulter). Data from all experiments were uniformly analyzed using FlowJo v10.8 (BD Biosciences).

### Single-cell V(D)J sequencing

For the 10X Genomics workflow, sorted antigen-specific B cells, suspended in PBS with 10% (v/v) FBS, were processed with the Chromium Next GEM Single Cell V(D)J Reagent Kits v1.1 (10X Genomics). The cell suspension was loaded onto a 10X Chromium Controller to generate Gel Beads-in-Emulsion (GEMs), which facilitate the barcoding of mRNA and subsequent reverse transcription within individual droplets. Following cDNA synthesis, the product was purified using a SPRIselect Reagent Kit (Beckman Coulter) and pre-amplified. Targeted enrichment of paired V(D)J sequences was then achieved using 10X-specific BCR primers, and the resulting products were used for sequencing library construction. These final libraries were sequenced on an Illumina NovaSeq 6000 platform with a NovaSeq 6000 S4 Reagent Kit v1.5 (Illumina).

### HTGTS-Rep-seq

0.5-4 µg of genomic DNA from purified splenic B cells was used for generating HTGTS-Rep-seq libraries as previously described ^54^. Four bait primers that target mouse JH1, JH2, JH3, and JH4 were mixed to capture all heavy chain (HC) repertoire in one library. All primers carried a 5′ biotin modification (5BiosG). These HTGTS-Rep-seq libraries were sequenced on DNBSEQ-T7.

### Monoclonal antibody expression and purification

The sequences for the antibody heavy and light chains were initially codon-optimized for expression in human cells (GenScript). The variable regions (VH and VL) were then separately inserted into corresponding expression vectors (pCMV3-CH, pCMV3-CL or pCMV3-CK). Plasmids of the heavy and light chain constructs were transformed into *Escherichia coli* DH5α competent cells (Tsingke). After overnight incubation at 37°C, single colonies were picked for colony PCR identification. Plasmid DNA of expanded cultures was extracted (CWBIO) after verified by Sanger sequencing.

For protein production, heavy and light chain plasmids were co-transfected into Expi293F cells using polyethylenimine (PEI; Yeasen). The plasmid-PEI complexes were prepared in sterile 0.9% NaCl solution before being added to the cell culture. The transfected cells were cultured at 37°C with 5% CO₂ and 175 rpm shaking for 6–10 days. A nutrient supplement (OPM Biosciences) was added to each culture at 24 hours post-transfection and every 48 hours thereafter.

To purify the antibodies, the culture supernatant was first clarified by centrifugation (3,000 × g, 10 minutes). The supernatant was then incubated with Protein A magnetic beads (GenScript) for 2 hours to allow antibody binding. The beads were subsequently washed, and the bound antibodies were eluted using a KingFisher automated purification system (Thermo Fisher). The concentration of the purified antibody was determined using a NanoDrop spectrophotometer (Thermo Fisher), and its purity was assessed by SDS-PAGE (LabLead).

### RBD depletion of serum

To deplete RBD-specific antibodies from serum, 50 μL of Dynabeads™ MyOne™ Streptavidin T1 (Invitrogen) were washed once with PBS. 10 μg of biotinylated SARS-CoV-2 Wuhan RBD was incubated with the washed beads for 1 hours with gentle rotation to allow binding via streptavidin-biotin interaction. The beads were then collected using a magnetic rack for 2-3 min, the supernatant was discarded, and the beads were washed three times with PBS to remove unbound proteins. Subsequently, 200-400 μL of serum was incubated with the RBD-conjugated beads for 1 hours with gentle rotation to allow specific antibody binding. Finally, the tubes were placed on the magnetic rack, and the supernatant representing the RBD-depleted serum was carefully collected for downstream analyses.

### Enzyme-linked immunosorbent assays

High-binding 96-well plates were coated overnight at 4°C with SARS-CoV-2 Wuhan or BA.5 RBD proteins. The following day, plates were washed three times with 1×PBST and blocked with 250 μL 3–5% bovine serum albumin in 1×PBST for 2 hours at 37°C to prevent non-specific binding. After three additional washes, 100 µL of serially diluted antibodies or serum samples were added to the wells and incubated for 30 minutes at room temperature. Unbound antibodies were removed by five washes with 1×PBST. Subsequently, 100 µL of HRP-conjugated Goat anti-Mouse IgG (H+L) Cross-Adsorbed Secondary Antibody (Invitrogen) or Peroxidase AffiniPure Goat Anti-Human IgG (H+L) (Jackson Immunoresearch) was added and incubated for 30 minutes at room temperature. Following a final five washes, the signal was developed by adding 100 µL of TMB substrate (Solarbio) to each well and incubating for 8 minutes in the dark. The reaction was terminated by adding 50 µL of stop solution (Solarbio). The optical density (OD) was measured at 450 nm with a reference wavelength of 630 nm using a Multiskan FC microplate reader (Thermo Scientific). Final absorbance values were obtained by subtracting the OD630 reading from the OD450 reading for each well.

### Surface plasmon resonance

Surface plasmon resonance (SPR) experiments were conducted using a Biacore 8K+ system (Cytiva) at room temperature. For competitive binding assays, His-tagged SARS-CoV-2 BA.5 RBD protein (5 µg/mL) was immobilized onto an anti-His-tagged CM5 sensor chip (Cytiva) for 1 minute. Subsequently, a saturating concentration of the first antibody (Ab1, 20 µg/mL) was injected for 2 minutes, immediately followed by the injection of the second antibody (Ab2, 20 µg/mL) for another 2 minutes. The sensor surface was regenerated between cycles using a glycine solution (pH 1.5). All binding data were recorded and processed using Biacore 8K Evaluation Software (v4.0.8.20368). The degree of epitope competition was calculated using the following formula:

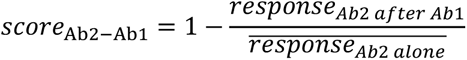

Where *response_Ab2 after Ab1_* represents the response units when Ab2 serves as the second antibody and Ab1 as the first antibody, whereas *response_Ab2alone_* denotes the mean response units when Ab2 acts as the first antibody.

### DMS library construction

Replicate deep mutational scanning (DMS) libraries of the SARS-CoV-2 BA.5 RBD (residues N331–T531; Wuhan-Hu-1 numbering) were generated based on established protocols ^14,24^ with a modification. Rather than using a pooled primer mix, we performed 201 individual PCR reactions, each using a specific NNS primer pair to introduce all possible amino acid substitutions at a single target residue. The resulting products from each single-site mutagenesis reaction were then combined to form the final comprehensive library. Each unique RBD variant was subsequently tagged with a distinct 26-nucleotide (N26) barcode via PCR. The mutagenized and barcoded RBD sequences were then cloned into pETcon 2649 vector and the resulted plasmid libraries were amplified in electrocompetent *E. coli* DH10B cells. The association between every RBD variant and its corresponding N26 barcode was established by preparing PacBio sequencing libraries and performing long-read sequencing on the Sequel II platform. Amplified DMS plasmid libraries were transformed into *Saccharomyces cerevisiae* strain EBY100. Transformed yeast cells were initially selected on SD-CAA agar plates. Positive clones were then expanded by culturing in SD-CAA liquid media. The resulting comprehensive DMS yeast libraries were preserved by flash-freezing in liquid nitrogen and stored at -80 °C.

### Magnetic beads-based antibody mutation escape profiling

High-throughput mutation escape profiling for mAbs was performed using magnetic beads based on previously established protocols ^14,24^. DMS yeast libraries (Wuhan and BA.5) first underwent functional pre-screening. Non-functional or misfolded RBD variants were removed using ACE2-biotin conjugate (Sino Biological) bound to streptavidin magnetic beads (Thermo Fisher). ACE2-bound yeast cells were washed with PBS containing 0.1% (v/v) BSA, released, expanded in SD-CAA liquid medium, and cryopreserved at -80°C as functional libraries.

For Antibody Escape Selection, thawed functional libraries were cultured overnight in SD-CAA with shaking, then back-diluted into SG-CAA medium to induce RBD surface expression. Escape variants of each mAb were isolated using a sequential selection strategy, with 2 rounds of negative selection to deplete antibody-binding variants and 1 round of positive selection to capture the antibody-escaping population using anti-c-Myc magnetic beads (Thermo Fisher). The final sorted yeasts were washed, regrown overnight and subjected to plasmid extraction using a 96-well kit (Coolaber). The unique N26 barcodes appended to each RBD variant were amplified by PCR using extracted plasmid as template. PCR products were purified with Ampure XP beads (Beckman Coulter) and subjected to high-throughput single-end sequencing (NextSeq 500/550 platforms or MGI200 platforms).

### Quantification and statistical analysis

#### Statistical analysis

Details of specific statistical tests and experimental design are given in the relevant figure legends. Data from neutralization assays were analyzed using GraphPad Prism (v8.0 for mice data and v9.0.1 for others). FACS, neutralization and ELISA data are visualized by R package ggplot2 (v3.5.0) and Python package matplotlib (v3.8.4). Statistical analysis was performed with a two-tailed Wilcoxon rank-sum test or paired Wilcoxon signed-rank test. P values < 0.05 were considered significant (\**P* < 0.05, \*\**P* < 0.01, \*\*\**P* < 0.001, \*\*\*\**P* < 0.0001). Randomization and blinding were not performed as this is an observational study applying a uniform set of measurements across the panel of monoclonal antibodies and plasma, and the experiments were not designed to directly indicate any efficacy.

#### 10X V(D)J sequence analysis

The 10X Genomics V(D)J Illumina sequencing data were assembled as B cell receptor contigs and aligned to the B cell V(D)J reference using Cell Ranger (v.7.1.0) pipeline. For human-source IGH and IGK contigs, we use GRCh38 as reference. For mouse-source IGL contigs, we use GRCm38 as reference. Only the productive contigs and B cells with one heavy chain and one light chain were kept to remove doublets. The germline V(D)J genes were identified and annotated using IgBlast (v1.17.1) ^55^. SHM nucleotides and residues in the antibody variable domain were detected using Change-O toolkit (v.1.2.0) ^56^.

#### HTGTS-Rep-seq sequence analysis

The HTGTS-Rep-seq data were analyzed with the HTGTS-Rep-seq pipeline ^54^.

#### Antibody DMS data analysis

The raw sequencing data from the DMS were processed as previously described ^14^. Specifically, the barcode sequences detected from both the antibody-screened and reference libraries were aligned with a barcode-variant dictionary derived from PacBio sequencing data of the Wuhan and BA.5 DMS libraries using the alignparse (v.0.6.2) and dms_variants (v.1.4.3) tools. Ambiguous barcodes were excluded during the merging of yeast libraries. Only barcodes detected more than five times in the reference library were considered for further analysis. The escape score for a variant X, present in both the screened and reference libraries, was calculated as *F* × (*n*_X,ab_/*N*_ab_)/(*n*_X,ref_/*N*_ref_), where *F* is a scaling factor to normalize the scores to a 0–1 range, and *n* and *N* represent the numbers of detected barcodes for variant X and the total barcodes in the antibody-screened (ab) or reference (ref) samples, respectively. For antibodies subjected to DMS with multiple replicates using different mutant libraries, the final escape score for each mutation was averaged for subsequent analyses.

We used graph-based unsupervised clustering and embedding to assign an epitope group to each antibody and visualize them in a two-dimensional space. Initially, site escape scores (sum of mutation escape scores per residue) for each antibody were normalized to a sum of one, representing a distribution over RBD residues. The dissimilarity between two antibodies is defined based on the Pearson’s correlation coefficient of their escape score vectors using numpy (v1.25.2). A k-nearest-neighbour graph was constructed using the python-igraph module (v.0.9.6), and Leiden clustering was applied to assign a cluster to each antibody ^57^. Cluster names were manually annotated on the basis of the characteristic sites in the average escape profiles of each cluster, using the same nomenclature as our previously published DMS dataset ^14^. To visualize the dataset in two dimensions, uniform manifold approximation and projection was performed based on the k-nearest-neighbour graph using umap-learn module (v.0.5.2), and figures were generated using R package ggplot2 (v.3.3.3).

To compute the average immune pressure or identify escape hotspots using a collection of mAb DMS profiles, we incorporating two types of weight to account for the impact of each mutation on neutralizing activity and codon constraints at each residue. For codon usage constraints, mutations inaccessible through single nucleotide changes were assigned a weight of zero, whereas others received a weight of 1.0. We used Wuhan/D614G (Wuhan-Hu-1 reference genome) and BA.4/5 (EPI_ISL_11207535) to define one-nucleotide-accessible amino acid mutations. Neutralizing activity weights were calculated as −log10 (IC_50_), with IC_50_ values below 0.0005 or above 1.0 adjusted to 0.0005 or 1.0, respectively. Raw escape scores for each antibody were normalized by the maximum score across all mutants. The weighted score for each antibody and mutation was obtained by multiplying the normalized scores by the corresponding two weights, and the final mutation-specific weighted score was the sum of scores for all antibodies in the designated set, subsequently normalized to a 0–1 range. To visualize the calculated escape maps, sequence logos were generated using the Python module logomaker (v.0.8).

---

**Figure S1.**
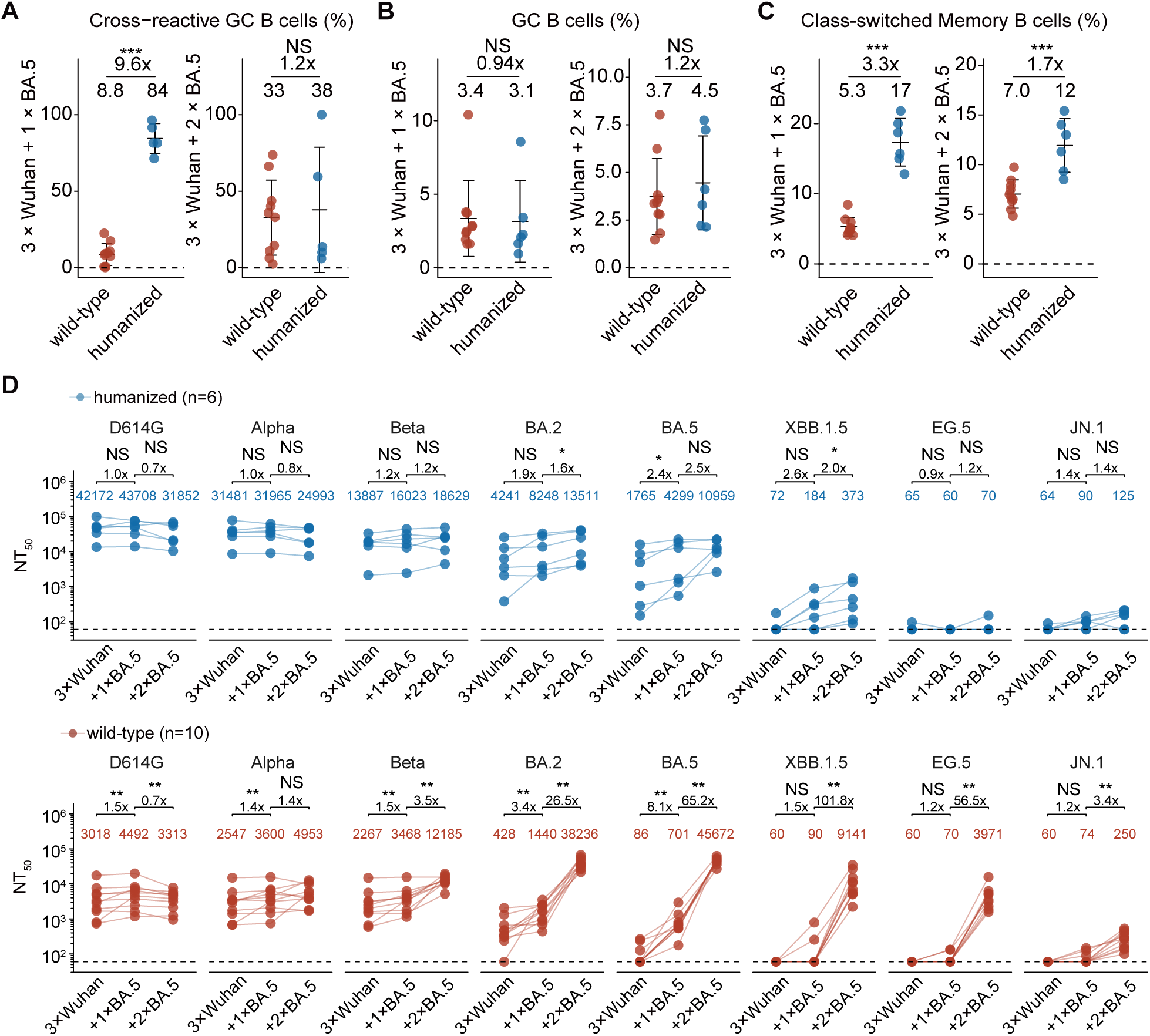
Cellular and serological characterization of SARS-CoV-2 imprinting in V(D)J-humanized mice, related to Figure 1. (A-C) Scatter plots showing the proportion of cross-reactive GC B cells (A), total GC B cells (B), and class-switched memory B cells (C). The indicated percentages reflect the frequency relative to their respective parental gates defined in Supplementary Information Figure 1A. Data are presented as mean ± standard deviation (SD). (D) Line plots showing longitudinal pseudovirus neutralization titres (NT_50_) against a panel of SARS-CoV-2 variants in paired humanized and wild-type mice across Wuhan priming and BA.5 booster timepoints; lines connect data from the same mouse. Dashed lines indicate the limit of detection (NT_50_ = 60). Paired Wilcoxon signed-rank tests were used in (D).

**Figure S2.**
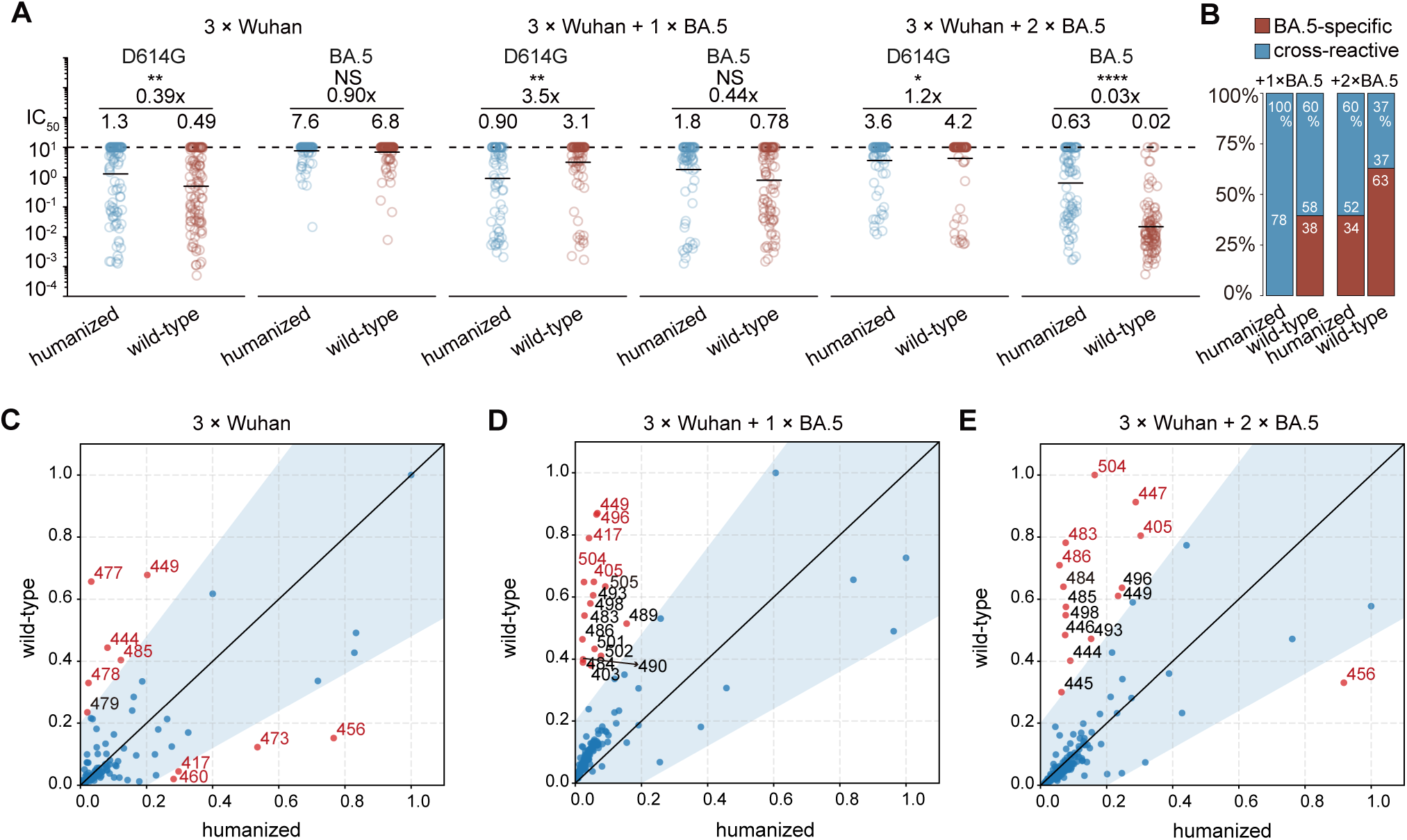
Comparative analysis of antibody potency and escape profiles across mouse strains, related to Figure 2. (A and B) Comparison of antibody IC50 values against D614G and BA.5 (A) and cross-reactivity proportion (B) from humanized and wild-type mice. Geometric mean values are displayed as bars and indicated above each group of data points in a. Two-tailed Wilcoxon rank-sum tests were used in a. Antibodies exhibiting ELISA OD450 values > 2 against both WT and BA.5 RBDs (1 μg/mL) were defined as cross-reactive. Those showing an OD450 > 2 for one variant but < 2 for the other were classified as specific. (C-E) Scatter plots comparing the normalized average DMS escape scores of neutralizing antibodies isolated from humanized (x-axis) versus wild-type (y-axis) mice following Wuhan priming (C), the first BA.5 booster (D), and the second BA.5 booster (E). Residues falling along the diagonal (y=x) indicate shared immune pressure between the two models. To identify divergent hotspots—including those with low-to-moderate scores in one strain that are absent in the other—a shaded tolerance region was defined by the boundaries y = 1.4x + 0.3 and y = 0.6x - 0.18. Points falling outside this shaded region represent distinct escape hotspots. Among these outliers, the five residues with the highest escape scores in each group are highlighted in red.

**Figure S3.**
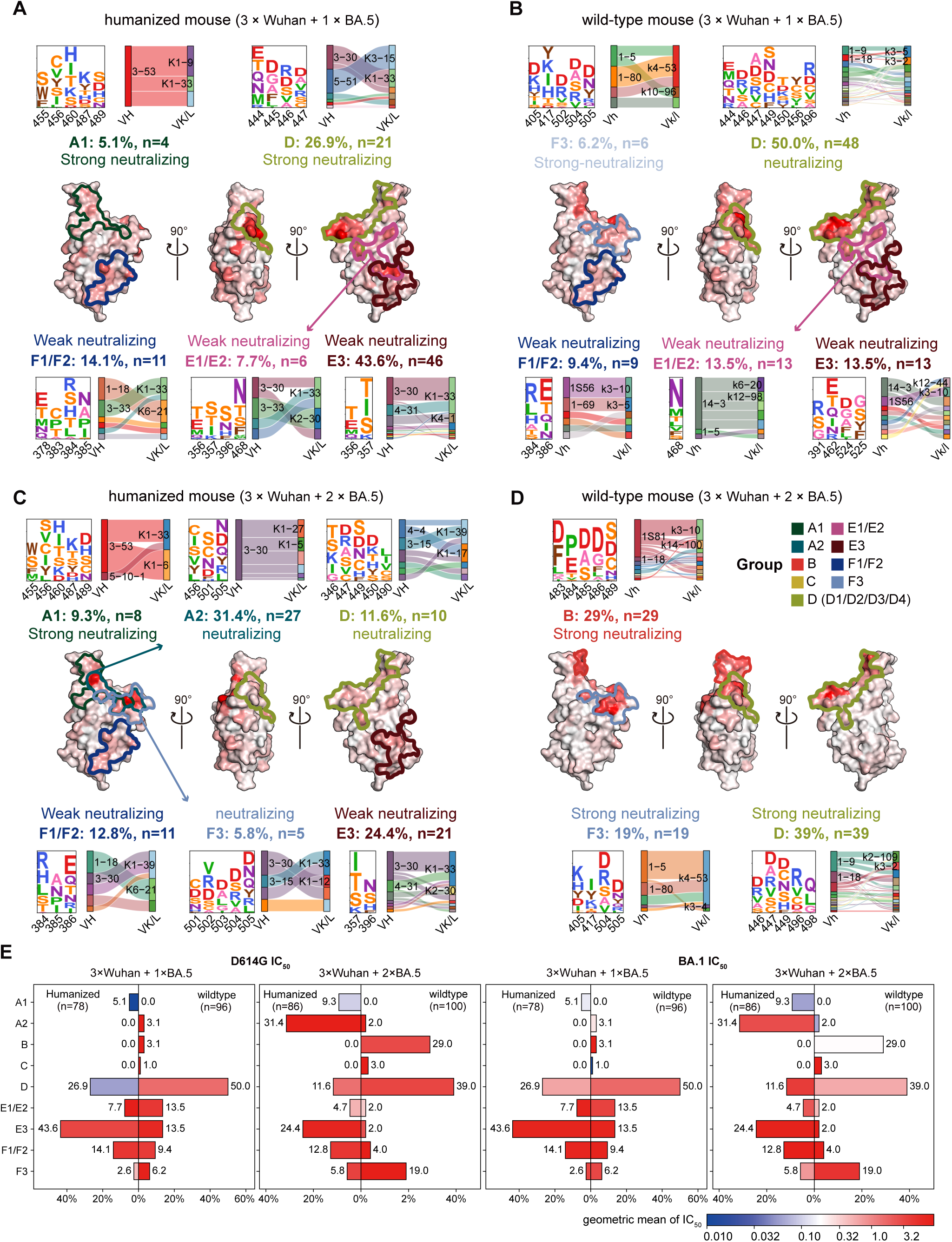
Distinct SARS-CoV-2 antibody epitope distribution after sequential BA.5 boosts, related to Figure 3. (A-D) Epitope distribution of the antibody repertoire generated after one BA.5 boost (A and B) or two BA.5 boosts (C and D) in humanized mice (A and C) and wild-type mice (B and D). E, Pyramidal bar charts showing the proportional distribution of epitope groups in antibodies isolated from humanized and wild-type mice after one or two BA.5 boosts. Bars are colored according to the log10 geometric mean IC_50_ of antibodies within each group.

**Figure S4.**
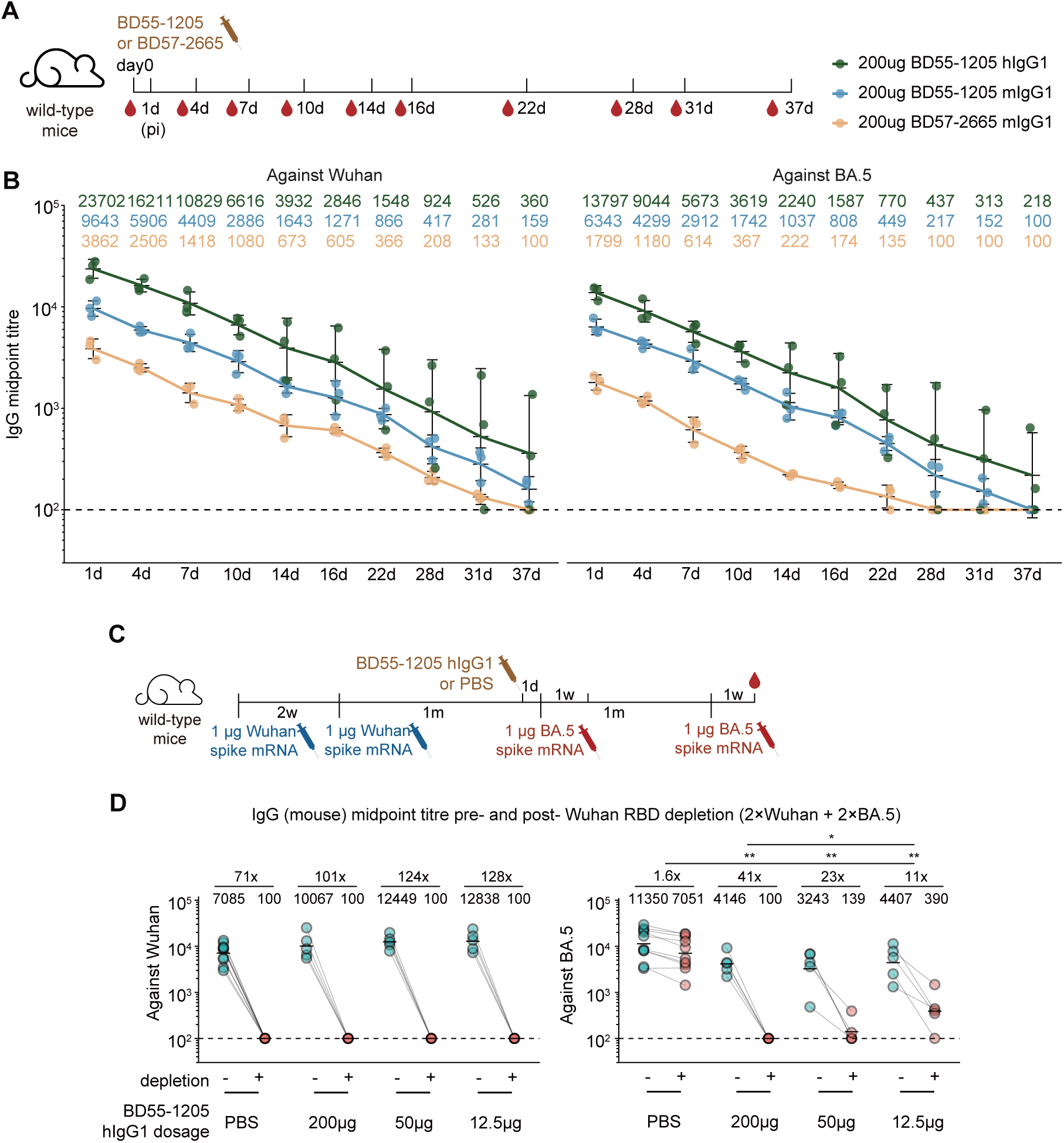
Pharmacokinetics and dose-dependent suppressive effects of antibodies used in passive transfer, related to Figure 4. (A) Schematic of the in vivo monoclonal antibody pharmacokinetic study. (B) Serum IgG titres over time following antibody injection (mIgG1 forms were detected using an anti-mouse Fc secondary antibody, and hIgG1 forms using an anti-human Fc secondary antibody). Dashed lines indicate the limit of detection (midpoint titre = 100). Data are presented as geometric mean titres (GMT), with error bars indicating geometric standard deviation. (C) Schematic of the BD55-1205 dose-ranging experiment. (D) Corresponding serum IgG titres before and after Wuhan RBD depletion. Geometric mean values are displayed as bars and indicated above each group of data points. Statistical significance of the fold-reduction in titres was assessed between groups. Dashed lines indicate the limit of detection (midpoint titre = 100). Two-tailed Wilcoxon rank-sum tests were used in (D).

**Figure S5.**
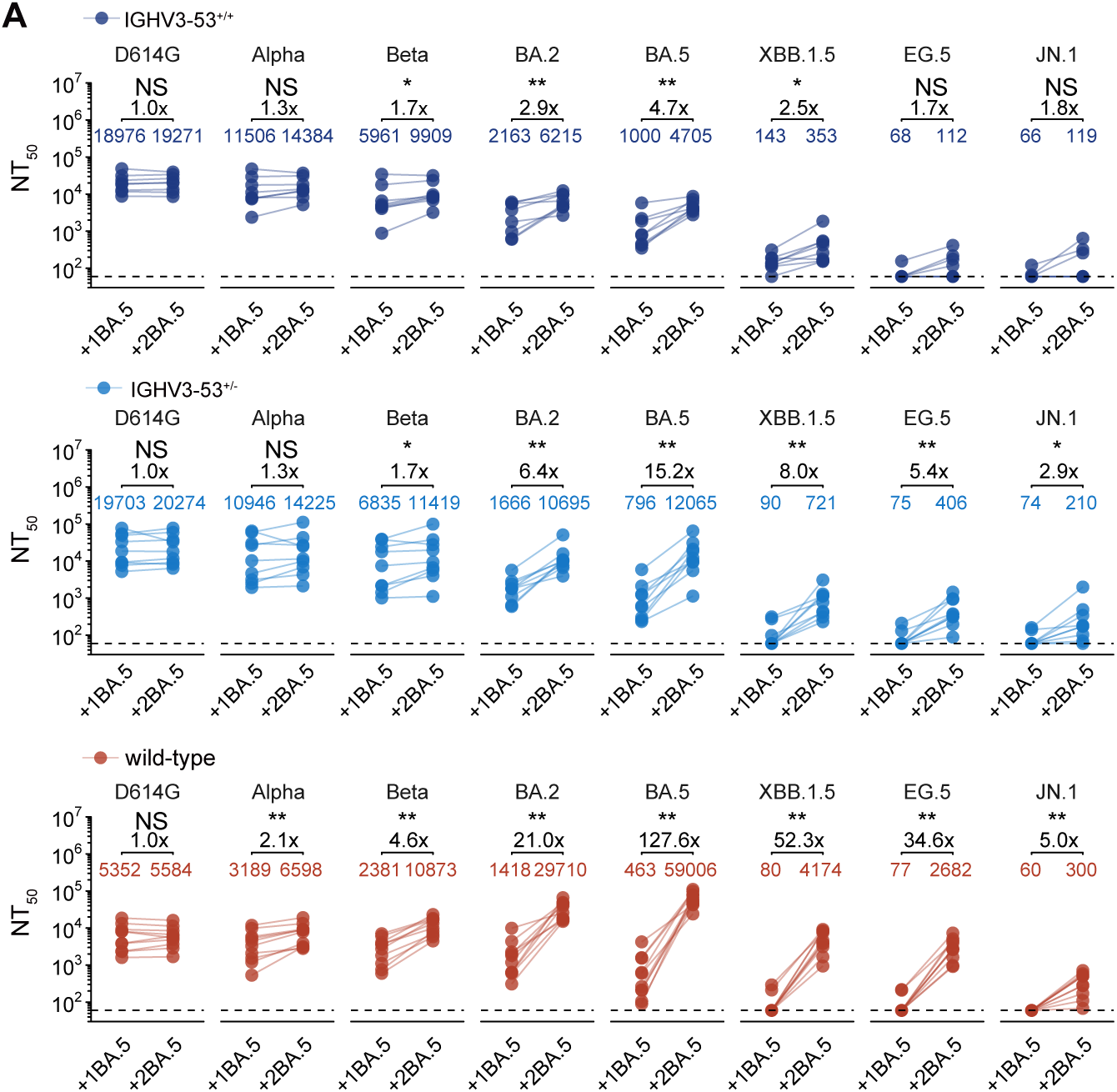
IGHV3-53 knock-in mice exhibit diminished Omicron neutralization boosting compared to wild-type mice, related to Figure 5. (A) Line plots showing longitudinal pseudovirus neutralization titres (NT_50_) against a panel of SARS-CoV-2 variants in paired IGHV3-53^+/+^ (top), IGHV3-53^+/-^ (middle), and wild-type (bottom) mice across BA.5 booster timepoints; lines connect data from the same mouse. Dashed lines indicate the limit of detection (NT_50_ = 60). Paired Wilcoxon signed-rank test were used.

**Figure S6.**
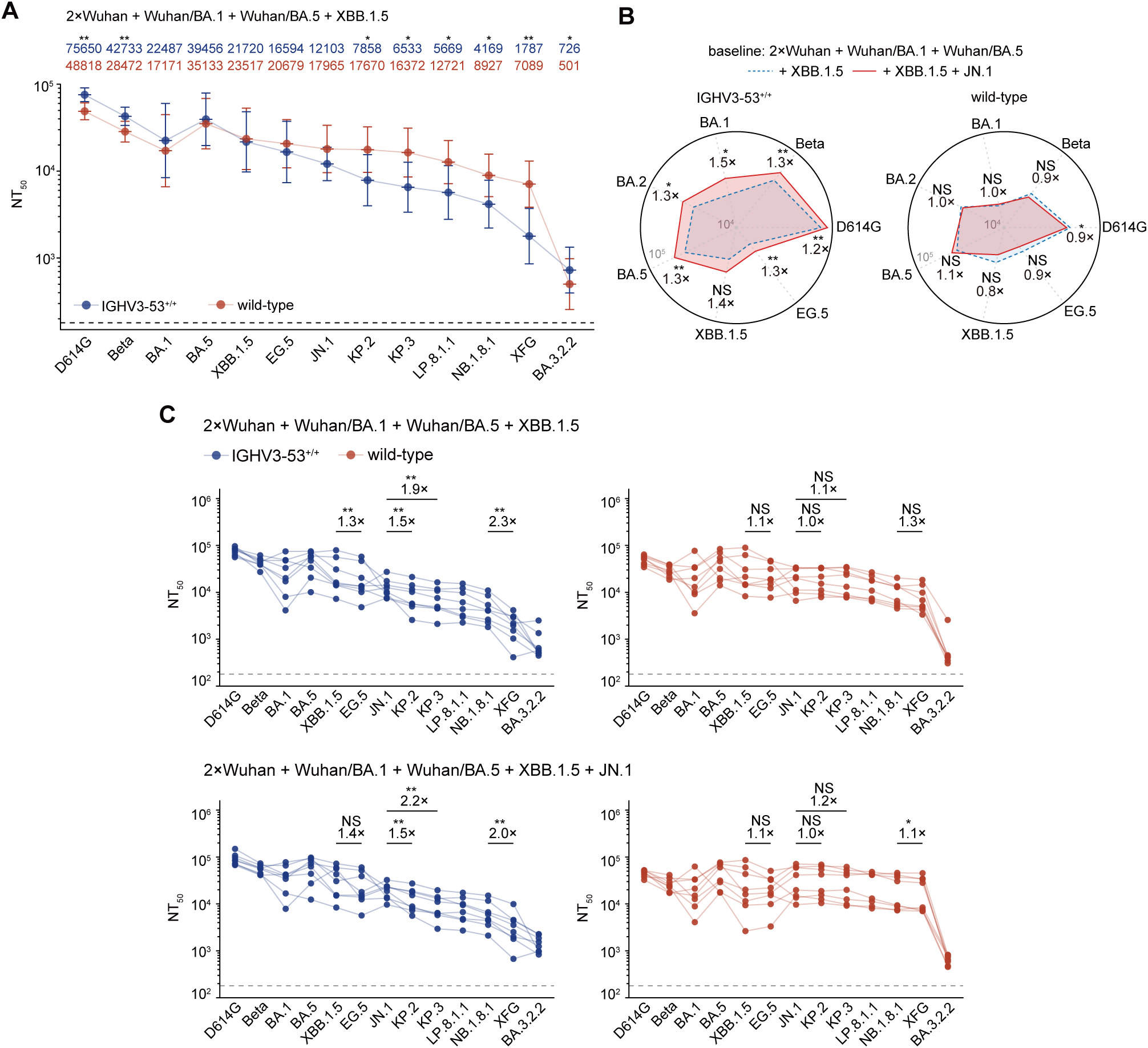
Strong back-boosting of Wuhan immunity in IGHV3-53 KI mice restricts the breadth of neutralization against emerging variants, related to Figure 6. (A) Serum neutralization titres (NT_50_) of IGHV3-53^+/+^ and wild-type mice against a panel of SARS-CoV-2 variant pseudoviruses following the fifth dose (XBB.1.5). Geometric mean titres (GMTs) are shown on the top. Dashed lines indicate the limit of detection (NT_50_ = 180). Data are presented as geometric mean titres (GMT), with error bars indicating geometric standard deviation. (B) Radar plot illustrating the back-boosting effect of the JN.1 booster on neutralization titres against pre-JN.1 variants. (C) Neutralization profiles of IGHV3-53^+/+^ and wild-type mice following XBB.1.5 and JN.1 boosters. Fold changes and statistical significance between highlighted variants are indicated. Lines connect data from the same mouse. Two-tailed Wilcoxon rank-sum tests were used in (A-B). Paired Wilcoxon signed-rank test were used in (C)

