## Supplementary Information for "Mouse V(D)J Humanization Recapitulates human-like Severe SARS-CoV-2 Immune Imprinting"

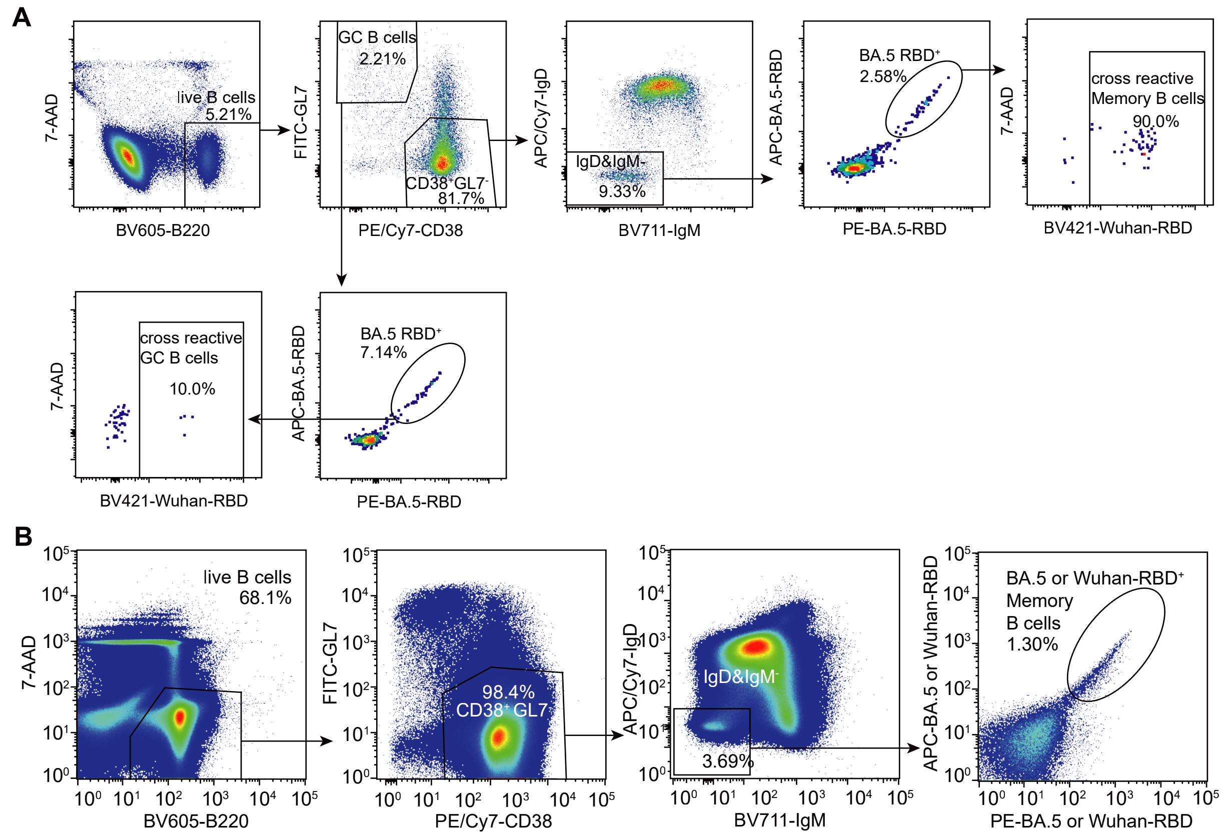


Supplementary Information Figure 1 FACS gating strategies.

**(A)** Representative gating strategy for FACS analysis of cross-reactivity in mouse memory B cells and GC B cells. APC/Cy7, APC/Cyanine7; BV711, Brilliant Violet 711. FITC, fluorescein isothiocyanate; BV605, Brilliant Violet 605; PE/Cy7, PE/Cyanine7.

**(B)** Representative gating strategy for sorting mouse Memory B cells.

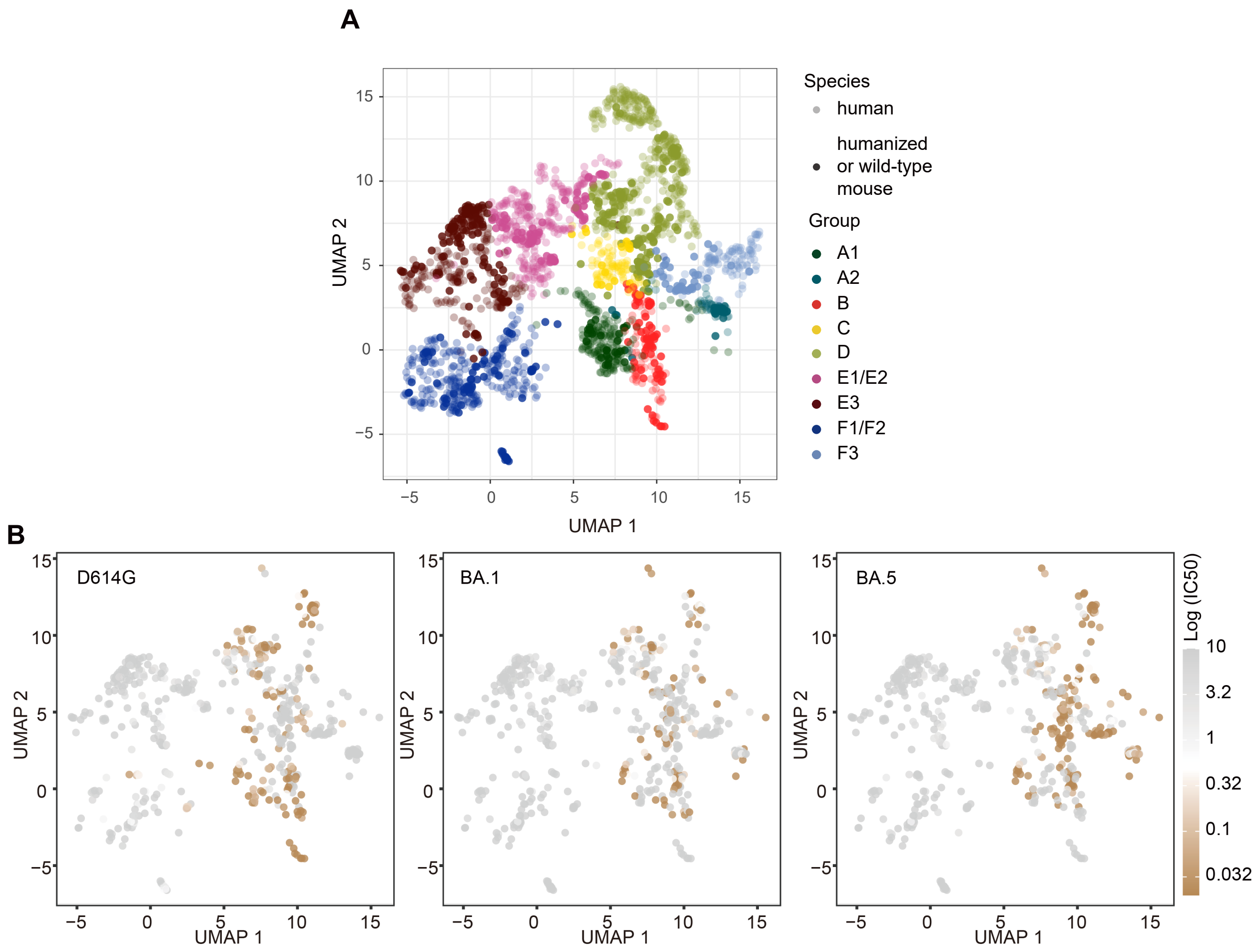


Supplementary Information Figure 2 UMAP-based epitope classification
**(A)** Uniform manifold approximation and projection (UMAP) visualization of antibody DMS escape mutation profiles. mAbs derived from wild-type and humanized mice in this study are highlighted with higher opacity, whereas reference human antibodies included to facilitate epitope clustering are displayed with lower opacity.
**(B)** UMAP visualization colored by IC50 values against D614G (left), BA.1 (middle), and BA.5 (right).

**A**

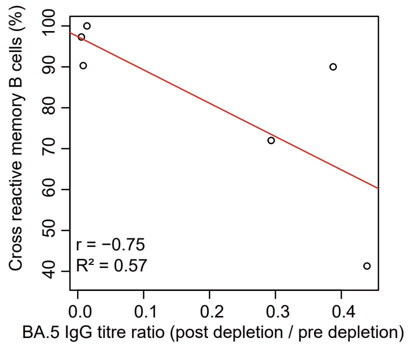


Supplementary Information Figure 3 Individual heterogeneity in imprinting strength of humanized mice

**(A)** Linear regression analysis of the correlation between memory B cell cross-reactivity proportion and BA.5-specific IgG titre. Simple linear regression was performed. The regression line (red) was fitted using the ordinary least squares (OLS) method. The Pearson correlation coefficient (r) and the coefficient of determination ($R^2$) were calculated to assess the strength of the linear relationship and the goodness of fit, respectively.

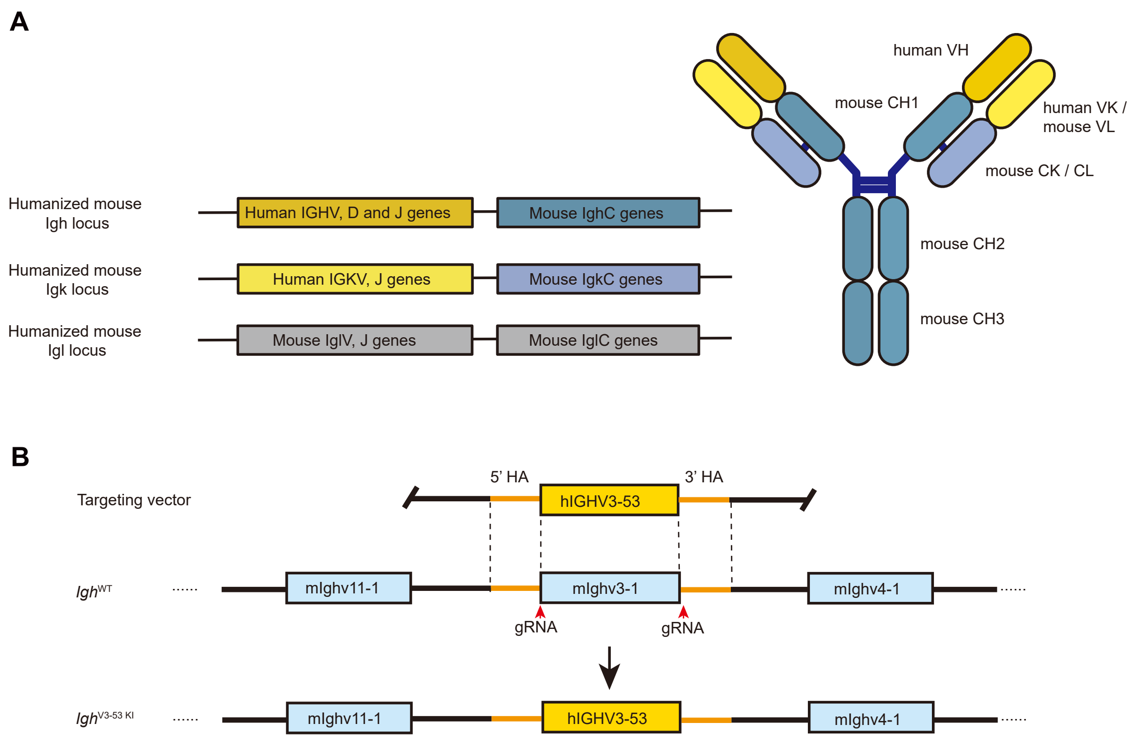


Supplementary Information Figure 4 immunoglobulin loci structure in VDJ-humanized and IGHV3-53 knock-in mice

**(A)** Schematic representation of the immunoglobulin loci in VDJ humanized mice. The variable regions of the heavy chain and kappa light chain are fully human, whereas the lambda light chain variable region and the constant regions of all chains remain murine.

**(B)** Schematic strategy for generating IGHV3-53 knock-in mice. The endogenous mouse Ighv3-1 gene segment was targeted and replaced by the human IGHV3-53 gene using CRISPR/Cas9-mediated genome editing. HA, homology arm.
